# Anle138b ameliorates pathological phenotypes in mouse and cellular models of Huntington’s disease

**DOI:** 10.1101/2025.03.11.642540

**Authors:** Miguel da Silva Padilha, Seda Koyuncu, Evangeline Chabanis, Sergey Ryazanov, Andrei Leonov, David Vilchez, Rüdiger Klein, Armin Giese, Christian Griesinger, Irina Dudanova

## Abstract

Huntington’s disease (HD) is a debilitating hereditary movement disorder caused by a CAG repeat expansion in the *huntingtin* gene. HD is characterized by deposition of mutant huntingtin (mHTT) aggregates, and by severe neurodegeneration of the basal ganglia and neocortex. No cure is currently available, and new treatment options are urgently needed. Here, we show that the oligomer modifying molecule anle138b (INN: emrusolmin) improves multiple disease phenotypes in cell culture and in two mouse models of HD. Application of anle138b reduced mHTT aggregate formation and ameliorated neurotoxicity in primary neurons. Oral administration of anle138b delayed deposition of mHTT inclusions, reduced brain atrophy, mitigated neuroinflammation, improved motor function and extended life span in HD mice. Downregulation of striatal markers and synapse loss in striatal spiny projection neurons were also partially rescued. No adverse effects of anle138b were observed in wildtype animals. Moreover, anle138b markedly decreased mHTT aggregation in human neural precursor cells differentiated from HD patient-derived induced pluripotent stem cells (iPSCs). Altogether these results illustrate the potential of anle138b as a disease-modifying treatment for HD.

## Introduction

Huntington’s disease (HD) is a hereditary brain disorder characterized by severe neurodegeneration, which affects in particular the basal ganglia and the neocortex (Waldvogel et al., 2015). HD manifests with motor defects, including involuntary movements (chorea) at early stages and akinesia at later stages, as well as psychiatric symptoms and cognitive impairments (Tabrizi et al., 2020). The cause of the disease is a CAG repeat expansion in exon 1 of the *huntingtin* (*HTT*) gene (The Huntington’s Disease Collaborative Research Group, 1993). The mutation is translated into a pathologically elongated polyglutamine (polyQ) tract in the HTT protein, resulting in its misfolding, aggregation, and deposition of mutant HTT (mHTT) inclusions in the brain (DiFiglia et al., 1997). HD leads to neuronal damage at least partially through the toxic gain of function of the aggregation-prone mHTT, in particular in the form of toxic oligomeric species. mHTT aggregation compromises cellular fitness through a variety of mechanisms including disturbance of protein homeostasis (proteostasis) (Labbadia and Morimoto, 2013; Miller et al., 2011; Saudou and Humbert, 2016; Tabrizi et al., 2020). Despite encouraging recent progress in the development of HTT-lowering therapies (Tabrizi et al., 2019), no cure for HD is yet available, and there is a great need for additional treatment options. Reducing mHTT aggregation might be a promising disease-modifying strategy.

The diphenylpyrazole compound anle138b [3-(1,3-benzodioxol-5-yl)-5-(3-bromophenyl)-1*H*-pyrazole] (INN: emrusolmin) was identified in a screen for small molecules with prion aggregation-inhibiting activity (Wagner et al., 2013). Anle138b was shown to directly interact with the oligomeric species of disease-related aggregating proteins such as prion protein, α-synuclein, amyloid-beta (Aβ), tau and human islet amyloid polypeptide, inhibiting formation of aggregates and reducing their toxicity (Albariqi et al., 2024; Heras-Garvin et al., 2019; Martinez Hernandez et al., 2018; Wagner et al., 2015; Wagner et al., 2013; Wegrzynowicz et al., 2019). The compound proved to have beneficial effects in mouse models of different neurodegenerative proteinopathies including prion disease, synucleinopathy, tauopathy and Aβ deposition (Brendel et al., 2019; Heras-Garvin et al., 2019; Levin et al., 2014; Martinez Hernandez et al., 2018; Wagner et al., 2015; Wagner et al., 2013). Anle138b is orally bioavailable, has suitable pharmacokinetic properties and efficiently penetrates the blood-brain barrier (Wagner et al., 2013). Importantly, a phase 1 clinical trial for anle138b demonstrated very good safety and tolerability profiles of the compound at exposure levels exceeding those required for full efficacy in mouse models (Levin et al., 2022). A phase 2 trial in multiple system atrophy patients is currently ongoing (NCT06568237). However, the therapeutic potential of anle138b in the context of HD has not been explored to date.

Here, we show that anle138b reduces mHTT aggregation in cultured primary neurons, two different mouse models of HD, as well as in human neural precursor cells (NPCs) differentiated from HD patients-derived induced pluripotent stem cells (iPSCs). Furthermore, anle138b mitigates neuropathological disease signatures, significantly improves motor performance, and extends life span of HD mice. These findings highlight the potential of anle138b as a treatment for HD.

## Results

### Anle138b reduces mHTT toxicity and aggregation in primary neurons

To assess the effects of anle138b in the context of HD, we first analyzed its effects on mHTT-dependent phenotypes in primary neuronal cultures isolated from mouse embryos. The neurons were transfected with two versions of mHTT-exon1, which differed in their polyQ length and tags: HTTQ72-His and HTTQ97-mCherry. We have shown in previous studies that these versions reproduce two characteristic locations of mHTT inclusion bodies: HTTQ72-His forms inclusions primarily in the nucleus, while inclusions of HTTQ97-mCherry are found mostly in the cytoplasm (Blumenstock et al., 2021; Voelkl et al., 2023). Immunostaining for the apoptotic marker cleaved caspase-3 was used to quantify neuronal cell death (Fig. 1A). Expression of both versions of mHTT significantly impaired neuronal survival (Fig. 1B), whereas the respective normal polyQ-length constructs (HTTQ25-His and HTTQ25-mCherry) did not cause neurotoxicity and were pooled qtogether as controls. Addition of anle138b to the culture medium resulted in a significant rescue of viability of both HTTQ72-His and HTTQ97-mCherry expressing neurons (Fig. 1B). In addition, anle138b reduced the fraction of neurons bearing mHTT inclusions for both versions of mHTT (Fig. 1C).

**Figure 1.**
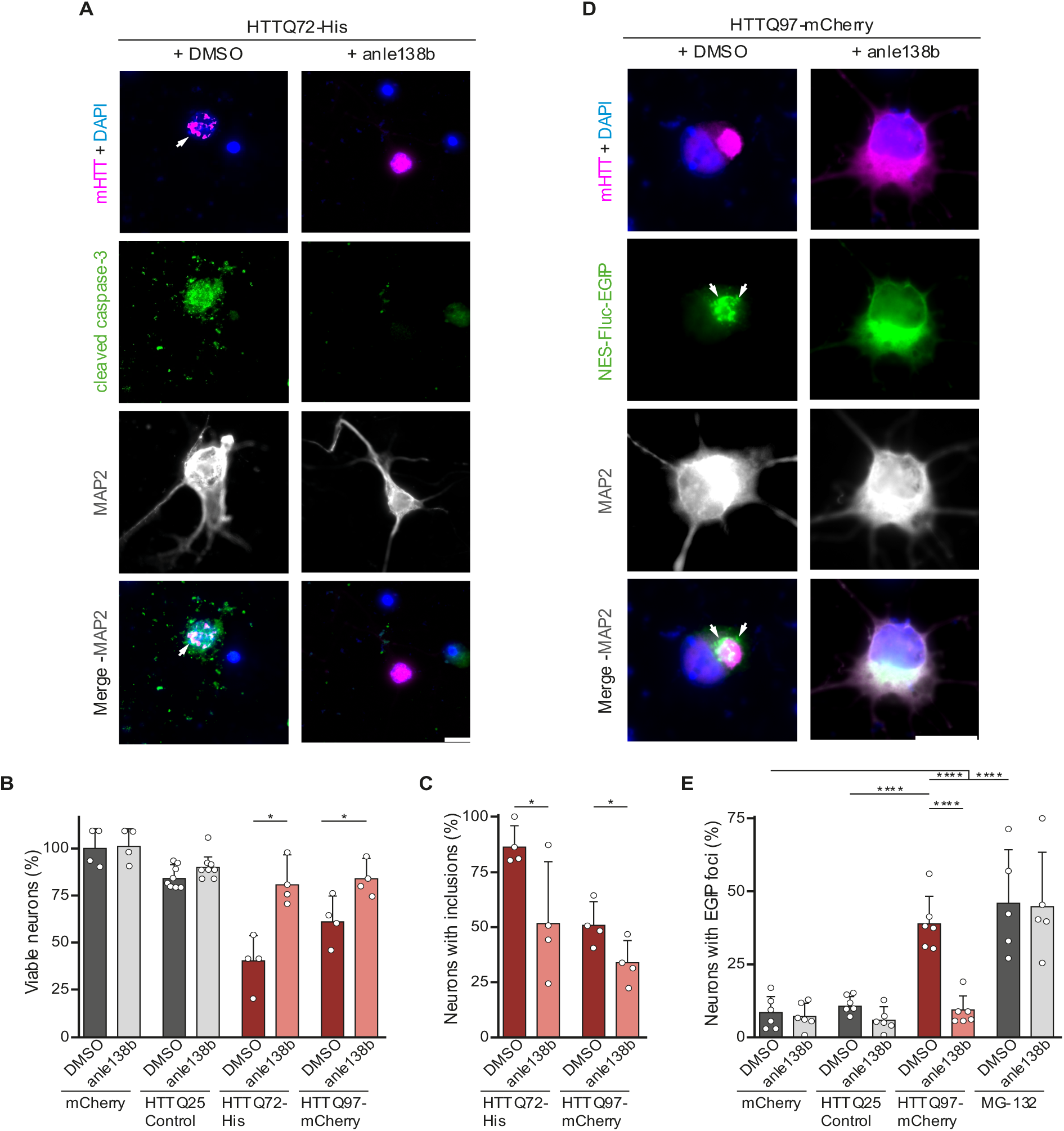
Anle138b reduces mHTT toxicity and improves proteostasis in primary neurons. **(A)** Representative images of DIV 7 + 2 cortical neurons transfected with HTT-Exon1-Q72-His and treated with vehicle (DMSO, left) or 7 µM anle138b (right). Neurons were stained for cleaved caspase-3 and neuronal marker MAP2, and mHTT was detected by immunostaining against the His-tag. Nuclei were labelled with DAPI. White arrows point to mHTT inclusion bodies. **(B)** Quantification of the percentage of viable transfected neurons, normalized to mCherry expressing cells treated with DMSO. Unpaired two-tailed *t-*test. n = 4 independent experiments. **(C)** Quantification of the fraction of neurons with mHTT inclusion bodies. Unpaired two-tailed *t-*test. n = 4 independent experiments. **(D)** Representative images of DIV 7 + 2 cortical neurons co-transfected with HTTQ97-mCherry and NES-Fluc-EGFP, treated with vehicle (DMSO, left) or 7 µM anle138b (right). Neurons were stained for MAP2, mHTT was detected by immunostaining against mCherry, and NES-Fluc-EGFP was detected by EGFP fluorescence. Nuclei were labelled with DAPI. White arrows point to EGFP foci. **(E)** Quantification of the fraction of double-transfected neurons with EGFP foci. Two-way ANOVA with Bonferroni’s multiple comparison test: Treatment, ****p < 0.0001; Construct, ****p < 0.0001; Treatment x Construct ***p = 0.001. n = 5 independent experiments for MG-132 and 6 independent experiments for all other conditions. Data are presented as mean ± SD. *p < 0.05, ****p < 0.0001. Scale bar in A and D, 10 µm.

To test whether the reduction in mHTT inclusion bodies is accompanied by improved neuronal proteostasis, we took advantage of the proteostasis reporter consisting of EGFP-fused firefly luciferase (Fluc-EGFP), which reveals proteostasis defects by forming EGFP foci in the cells (Gupta et al., 2011). We have previously shown that cytoplasmically targeted Fluc-EGFP (NES-Fluc-EGFP) reliably detects impairment in cytoplasmic proteostasis induced by HTTQ97-mCherry (Blumenstock et al., 2021). We therefore co-expressed NES-Fluc-EGFP and HTTQ97-mCherry in primary neurons and quantified EGFP foci upon treatment with anle138b or vehicle control (Fig. 1D). As a positive control, Fluc-EGFP transfected neurons were treated with the proteasome inhibitor MG-132, which readily induces Fluc-EGFP reaction. Consistent with our previous findings (Blumenstock et al., 2021), both HTTQ97-mCherry expression and proteasome inhibition, but not control HTTQ25-mCherry, caused a significant increase in the fraction of neurons with EGFP foci, indicative of impaired proteostasis. This impairment was completely rescued by anle138b application in the case of mHTT, but not MG-132 (Fig. 1E), suggesting that anle138b specifically rectifies proteostasis defects caused by aggregation of mHTT. Taken together, these results indicate that anle138b reduces aggregation of mHTT and prevents mHTT-dependent proteostasis failure and cell death in primary neurons.

### Anle138b improves motor function and extends life span in R6/2 mice

To explore the therapeutic potential of anle138b *in vivo*, we first used the R6/2 mouse line, which is a transgenic fragment model of HD that expresses exon 1 of human *HTT* with an expanded CAG tract under the human *HTT* promoter (Appendix Fig. S1A) (Mangiarini et al., 1996). The R6/2 model is well-characterized, has a rapid disease progression, clear histological and behavioral phenotypes and a shortened life span (Burgold et al., 2019; Carter et al., 1999; Mangiarini et al., 1996). Non-transgenic littermates were used as control throughout the study. Since HD is a monogenic disorder, and gene expansion carriers can be identified at an early age or even prenatally through genetic testing, we treated the mice from conception in order to achieve the maximal possible effect of anle138b. The compound was administered in chow at the concentration of 2 g/kg, and placebo food with the same composition but without anle138b served as control. Mice were tested in a battery of assays for motor behavior at the age of 8 weeks (around onset of motor impairments) and 12 weeks (advanced stage). Some of the animals were sacrificed after the behavioral tests and brains were taken out for histological and biochemical analyses, while the rest of the mice were used to measure life span (Fig. 2A).

**Figure 2.**
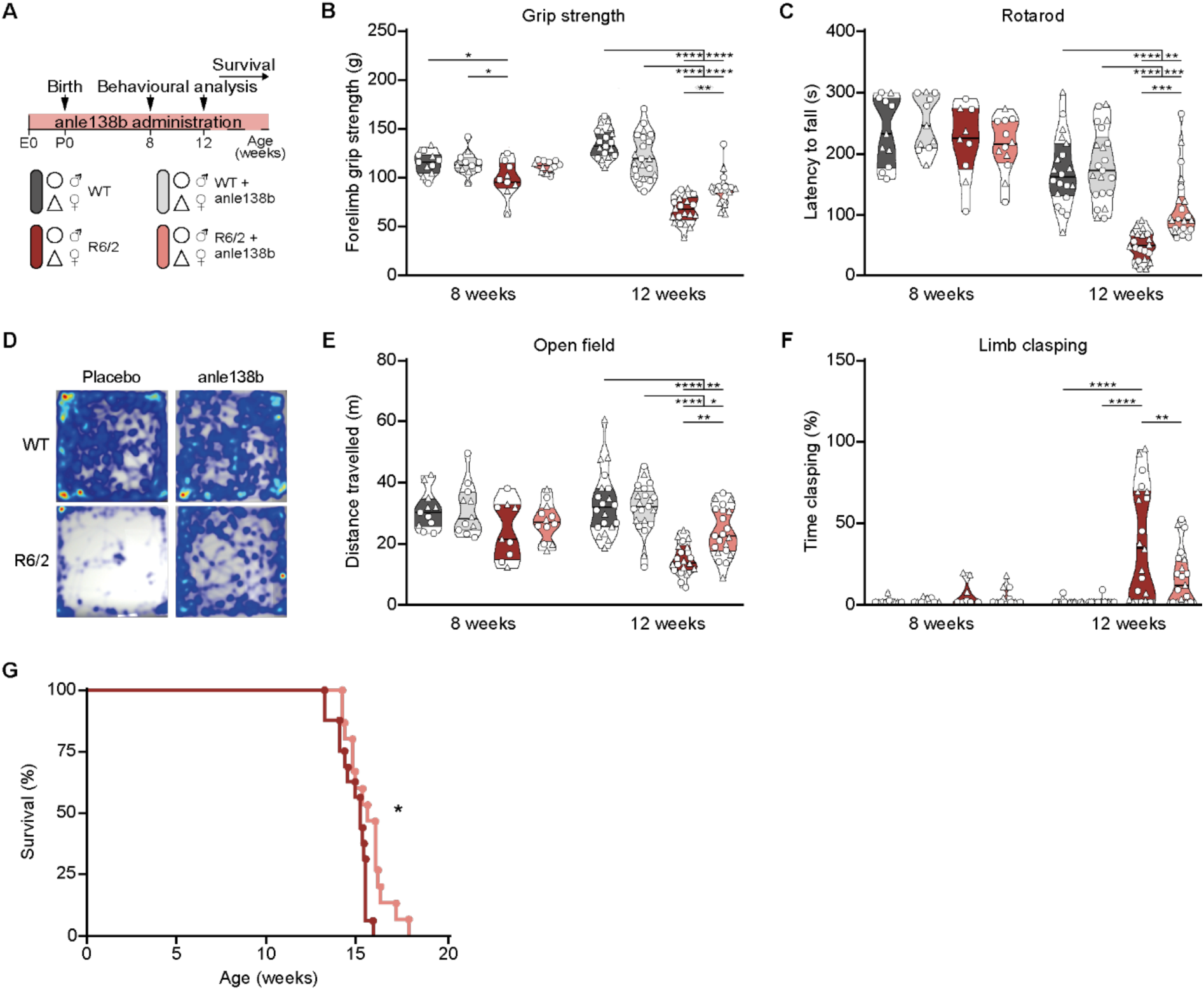
Anle138b improves motor performance and extends life span in R6/2 mice. **(A)** Experimental timeline for the R6/2 mice. **(B)** Forelimb grip strength. Two-way ANOVA with Bonferroni’s multiple comparison test, per age group. ANOVA for the 8 weeks’ time point: Treatment, p = 0.0801; Genotype, **p = 0.0073; Treatment x Genotype, p = 0.0839. ANOVA for the 12 weeks’ time point: Treatment, p = 0.3747; Genotype, ****p < 0.0001; Treatment x Genotype, ****p < 0.0001. **(C)** Latency to fall from the rotarod. Two-way ANOVA with Bonferroni’s multiple comparison test, per age group. 8 weeks: not significant; 12 weeks: Treatment, **p = 0.0023; Genotype, ****p < 0.0001; Treatment x Genotype, *p = 0.0141. **(D)** Heatmap of the open field arena depicting preferred occupied locations, from less (dark blue) to more (red) time spent in the same location. **(E)** Distance travelled in the open field arena. Two-way ANOVA with Bonferroni’s multiple comparison test, per age group. 8 weeks: not significant; 12 weeks: Treatment, *p = 0.0342; Genotype, ****p < 0.0001; Treatment x Genotype, **p = 0.0053. **(F)** Fraction of time spent clasping. Two-way ANOVA with Bonferroni’s multiple comparison test, per age group. 8 weeks: not significant; 12 weeks: Treatment, *p = 0.0131; Genotype, ****p < 0.0001; Treatment x Genotype, *p = 0.0140. **(G)** Kaplan-Meier Survival curve. Log-rank test. For all behavioral analyses (B - F), n = 9 – 12 (8 weeks) or 19 – 22 mice (12 weeks) per genotype and treatment group. Males and females are represented by circles and triangles, respectively. Data presented as violin plots with median and interquartile ranges. Significant pairwise comparisons are indicated on the graphs. *p < 0.05, **p < 0.01, ***p < 0.001, ****p < 0.0001.

Anle138b treatment did not change the litter size (Appendix Fig. S1B) or the Mendelian distribution of genotypes, suggesting that it does not cause adverse effects on prenatal development. Placebo-treated R6/2 mice showed diminished forelimb strength in the grip strength test starting from 8 weeks. At 12 weeks, they also exhibited impaired coordination on the rotarod and reduced locomotion in the open field. In addition, when suspended by the tail, the mice clasped their limbs, a characteristic phenotype of HD animals (Fig. 2B-F). All these motor defects were significantly mitigated in anle138b-treated R6/2 mice. The compound did not have an effect on the body weight of the mice (Appendix Fig. 1C). Of note, no significant effect of anle138b on wildtype control littermates was observed in any of the tests (Fig. 2B-F and Appendix Fig. S1C). In addition, administration of anle138b prolonged the life span of R6/2 mice by 5 days on average (Fig. 2G). Analyses of the mice separated by sex showed that the difference in survival was due to a significantly longer life span of female mice, while the life span of the male mice was not changed (Appendix Fig. S1D-E). Altogether, these findings demonstrate that oral administration of anle138b ameliorates HD-related neurological phenotypes and increases the life span of R6/2 animals.

### Anle138b mitigates brain atrophy and astrogliosis in R6/2 mice

To investigate the impact of anle138b treatment on the neuropathological hallmarks of HD, we performed histological analysis of the R6/2 mouse brains at 8 and 13 weeks of age. Placebo-treated R6/2 mice had a significantly smaller forebrain at 13 weeks, a defect that was partially rescued by anle138b (Fig. 3A-B). To assess brain atrophy in more detail, we performed stereological analyses of serial coronal brain sections. The estimated total volume of the brain was significantly smaller in placebo-treated R6/2 mice compared to littermate controls, whereas no significant difference was detected between anle138b-treated R6/2 animals and controls (Fig. 3C-D). Moreover, placebo-treated R6/2 mice showed atrophy of the striatum and motor cortex, as well as a pronounced enlargement of the brain ventricles. While we did not detect a significant effect of anle138b on the size of the striatum or motor cortex (Appendix Fig. S2A), the ventricle enlargement was partially rescued by anle138b administration (Fig. 3C-D).

**Figure 3.**
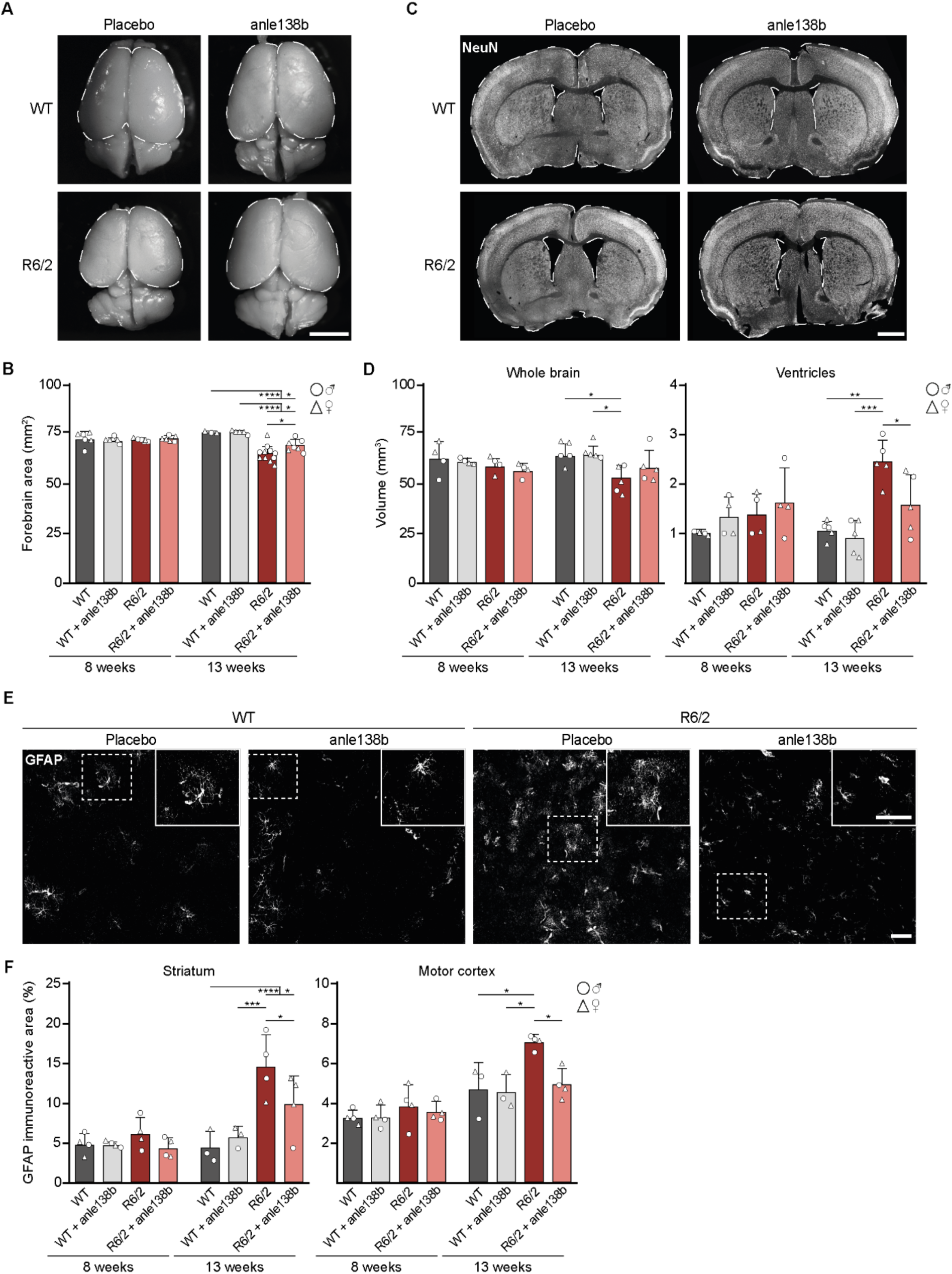
Treatment with anle138b ameliorates brain atrophy and decreases astrogliosis. **(A)** Representative whole brains of 13-week-old WT and R6/2 mice treated with placebo or anle138b. Dashed white lines outline the forebrain. **(B)** Quantification of whole forebrain area in 8 and 13-week-old WT and R6/2 mice. Two-way ANOVA with Bonferroni’s multiple comparison test, per age group. 8 weeks: not significant; 13 weeks: Treatment, p = 0.1260; Genotype, ****p < 0.0001; Treatment x Genotype, p = 0.1089. n = 6 (8 weeks) or 3 – 11 mice (13 weeks) per group. **(C)** Representative coronal brain sections stained with NeuN from 13-week-old WT and R6/2 mice treated with placebo or anle138b. Dashed white lines outline the whole brain and the ventricles. **(D)** Whole brain and ventricle volume quantification in 8 and 13-week-old WT and R6/2 mice. Two-way ANOVA with Bonferroni’s multiple comparison test, per age group and brain region. Whole brain, 8 weeks: not significant; 13 weeks: Treatment, p = 0.3006; Genotype, **p = 0.0047; Treatment x Genotype, p = 0.3168. Ventricle volume, 8 weeks: not significant; 13 weeks: Treatment, **p = 0.0052; Genotype, ****p < 0.0001; Treatment x Genotype, p = 0.1378. n = 4 (8 weeks) or 5 mice (13 weeks) per genotype and treatment group. **(E)** Representative astrocytes stained with GFAP in the dorsal striatum of 13-week-old WT and R6/2 mice treated with placebo or anle138b. Insets in the top right corner show magnification of the areas delineated by the dashed white lines. **(F)** Fraction of GFAP occupied area in the striatum (left) and motor cortex (right) of 8 and 13-week-old WT and R6/2 mice. Two-way ANOVA with Bonferroni’s multiple comparison test, per age group and brain region. 8 weeks: not significant; Striatum at 13 weeks: Treatment, ***p = 0.0003; Genotype, ***p = 0.0003; Treatment x Genotype, **p = 0.0041. Motor cortex at 13 weeks: Treatment, *p = 0.0328; Genotype, *p = 0.0128; Treatment x Genotype, p = 0.0561. n = 4 (8 weeks) or 3 – 4 mice (13 weeks). Data presented as mean ± SD. Significant pairwise comparisons are indicated on the graphs. *p < 0.05, **p < 0.01, ***p < 0.001, ****p < 0.0001. Scale bars: A, 5 mm; C, 1 mm; E, 100 µm.

A prominent feature of HD is neuroinflammation, characterized by altered morphology and density of astrocytes and microglial cells (Palpagama et al., 2019; Wilton and Stevens, 2020). To assess the impact of anle138b on astrogliosis and microgliosis, we performed immunostainings for the astrocyte marker glial fibrillary acidic protein (GFAP) and microglial marker Iba1, and quantified the immunoreactive area in the striatum and motor cortex. GFAP immunoreactive area was significantly increased in both brain regions of placebo-treated R6/2 mice at 13 weeks of age. This increase was diminished by anle138b treatment (Fig. 3E-F). We also observed a trend towards increased Iba1 immunoreactive area in placebo-treated R6/2 mice, but this effect did not reach statistical significance, precluding us from assessing the impact of anle138b on microglia (Appendix Fig. S2B-C). In summary, histological analyses revealed beneficial effects of anle138b on brain size and neuroinflammation markers in the R6/2 mouse brain.

### Anle138b reduces mHTT inclusion load and reverses neurochemical changes in the R6/2 brain

As anle138b reduced the frequency of mHTT inclusion bodies in cultured neurons, we next assessed its impact on mHTT inclusion load in the mouse brain. EM48 staining for aggregated HTT revealed abundant neuronal inclusions in the cortex and striatum of R6/2 mice (Fig. 4A), as described previously (Davies et al., 1997; Hosp et al., 2017). While anle138b did not have a pronounced effect on the inclusion size (Appendix Fig. S2D), it significantly reduced the fraction of inclusion-positive neurons (Fig. 4B), with the exception of 13-week-old striatal samples, an advanced time point when the immunohistochemically detectable inclusion load in the striatum is saturated. In line with the histological analyses, filter trap assay revealed a decrease in aggregated mHTT in anle138b-treated brains from 8 weeks on (Fig. 4C-D).

**Figure 4.**
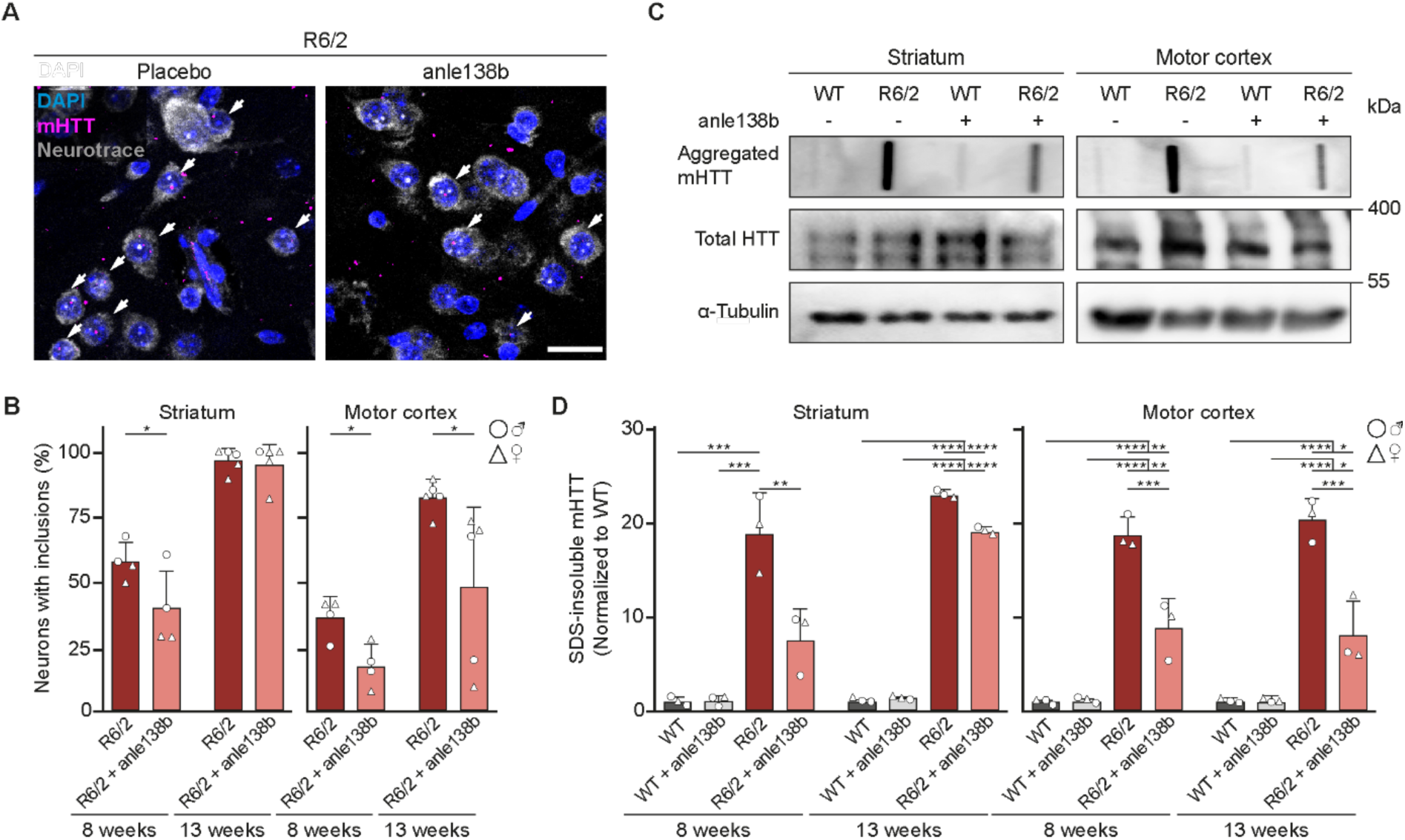
Anle138b reduces mHTT aggregate load in R6/2 mice. **(A)** Representative images of the motor cortex of 13-week-old R6/2 mice treated with placebo or anle138b. Neurons were identified by Neurotrace labeling and aggregated mHTT was detected by EM48 immunostaining. Nuclei were labelled with DAPI. White arrows point to neurons with mHTT inclusion bodies. **(B)** Quantification of the fraction of neurons with mHTT inclusion bodies in the striatum and motor cortex of R6/2 mice. Unpaired two-tailed *t* test. n = 4 (8 weeks) or 5 mice (13 weeks). **(C)** Filter trap membranes of lysates from the striatum and motor cortex of 8-week-old WT and R6/2 mice treated with placebo or anle138b. Total levels of HTT were determined by immunoblotting and α-Tubulin was used as a loading control. **(D)** Quantification of the filter trap assay. Values were normalized to WT/placebo. Two-way ANOVA with Bonferroni’s multiple comparison test. Striatum at 8 weeks: Treatment, **p = 0.0064; Genotype, ****p < 0.0001; Treatment x Genotype, **p = 0.0061. Striatum at 13 weeks: Treatment, ****p < 0.0001; Genotype, ****p < 0.0001; Treatment x Genotype, ****p < 0.0001. Motor cortex at 8 weeks: Treatment, **p = 0.0016; Genotype, ****p < 0.0001; Treatment x Genotype, **p = 0.0014. Motor cortex at 13 weeks: Treatment, **p = 0.0012; Genotype, ****p < 0.0001; Treatment x Genotype, ***p = 0.0007. n = 3 mice per group. Data presented as mean ± SD. Significant pairwise comparisons are indicated on the graphs. *p < 0.05, **p < 0.01, ***p < 0.001, ****p < 0.0001. Scale bar in A, 20 µm.

Neurodegenerative changes in the striatum of HD mice include the loss of medium spiny neuron markers such as dopamine- and cAMP-regulated neuronal phosphoprotein of 32 kDa (DARPP-32) and phosphodiesterase 10A (PDE10A) (Bibb et al., 2000; Menalled et al., 2012; Suelves et al., 2019). PDE10A downregulation also occurs in HD patients, and PDE10 tracers are used in human positron emission tomography (PET) imaging as an early-stage HD biomarker (Niccolini et al., 2015; Russell et al., 2014). We therefore quantified these marker proteins in the striatum of anle138b treated mice by western blot. R6/2 mice displayed reduced levels of both DARPP-32 and PDE10A in striatal lysates at 13 weeks of age. Remarkably, anle138b administration reversed the loss of both marker proteins (Fig. 5A-D). These results indicate that anle138b treatment decreases mHTT aggregate load and prevents loss of striatal neuron markers in R6/2 mice.

**Figure 5.**
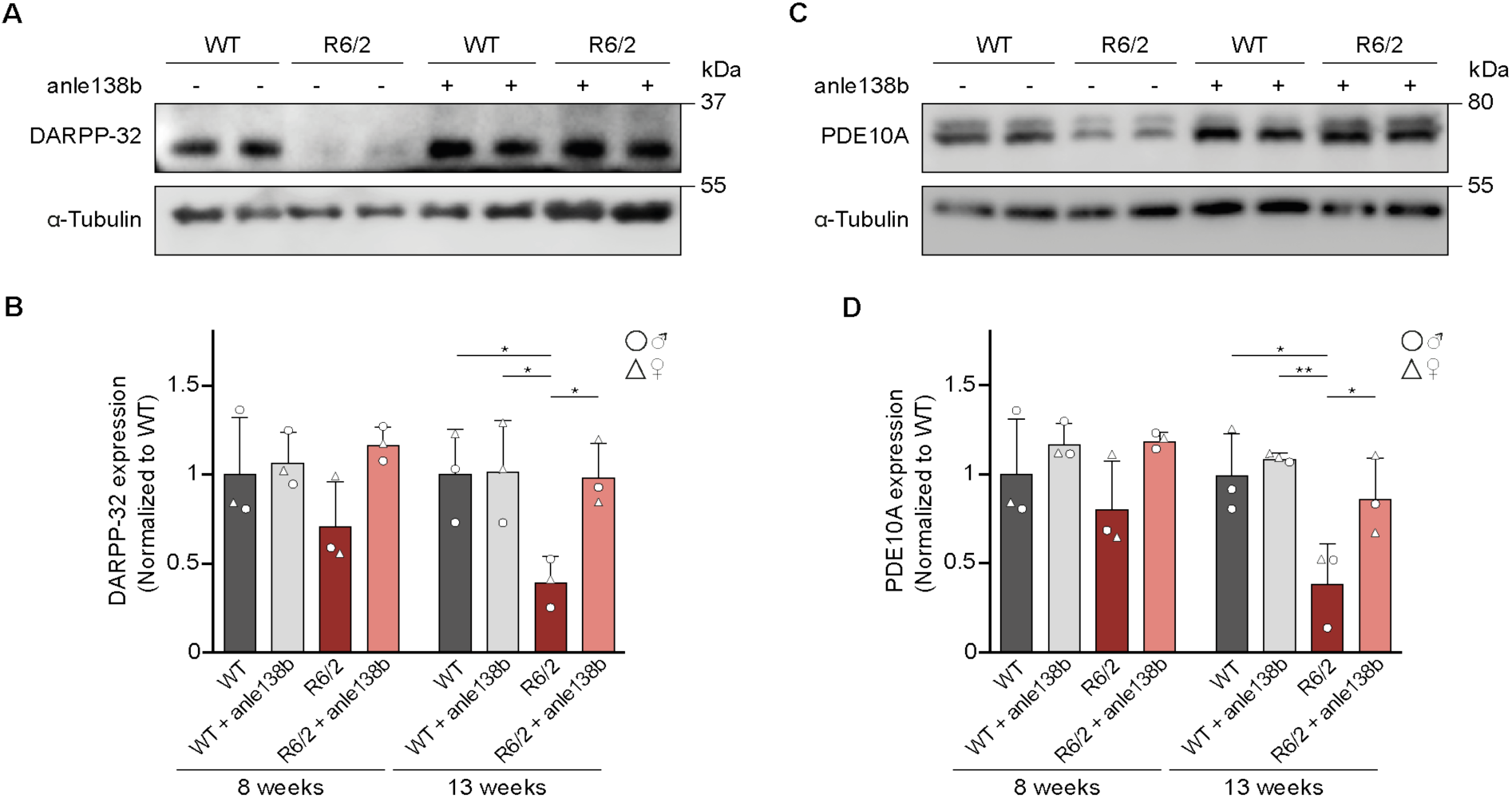
Anle138b treatment rescues expression of striatal markers in R6/2 mice. **(A)** Representative immunoblot for DARPP-32 in striatal lysates of 13-week-old WT and R6/2 mice treated with placebo or anle138b. **(B)** Quantification of DARPP-32 expression levels in the striatum of 8 and 13-week-old WT and R6/2 mice. Values were normalized to WT/placebo. Two-way ANOVA with Bonferroni’s multiple comparison test. 8 weeks: not significant; 13 weeks: Treatment, *p = 0.0386; Genotype, *p = 0.0222; Treatment x Genotype, *p = 0.0334. **(C)** Representative immunoblot for PDE10A in striatal lysates of 13-week-old WT and R6/2 mice treated with placebo or anle138b. **(D)** Quantification of PDE10A expression levels in the striatum of 8 and 13-week-old WT and R6/2 mice. Values were normalized to WT/placebo. Two-way ANOVA with Bonferroni’s multiple comparison test. 8 weeks: not significant; 13 weeks: Treatment, *p = 0.0215; Genotype, **p = 0.0035; Treatment x Genotype, p = 0.0711. In A and C, α-Tubulin was used as loading control. For all assays, n = 3 mice per group. Data presented as mean ± SD. Significant pairwise comparisons are indicated on the graphs. *p < 0.05, **p < 0.01.

### Anle138b mitigates synapse loss in the striatum of R6/2 mice

Synaptic defects constitute an important hallmark of neurodegenerative diseases including HD, where loss of glutamatergic corticostriatal inputs precedes and likely triggers dysfunction and degeneration of medium spiny neurons (Deng et al., 2013; Estrada-Sanchez and Rebec, 2013; Langfelder et al., 2016; Spampanato et al., 2008; Uytterhoeven et al., 2025). Previous studies demonstrated reduction in dendritic spines and downregulation of synaptic marker proteins in HD mouse models (Hosp et al., 2017; Indersmitten et al., 2015; Langfelder et al., 2016; Morton and Edwardson, 2001; Murmu et al., 2013). We therefore investigated the ability of anle138b to ameliorate these impairments in the striatum of R6/2 mice. First, we visualized the dendritic arbors of the striatal medium spiny neurons in brain sections by DiIC18 labeling and counted spines on the proximal dendrites. Dendritic spine density was significantly reduced in placebo-treated R6/2 mice at 13 weeks of age. This reduction was rescued by anle138b treatment (Fig. 6A-B).

**Figure 6.**
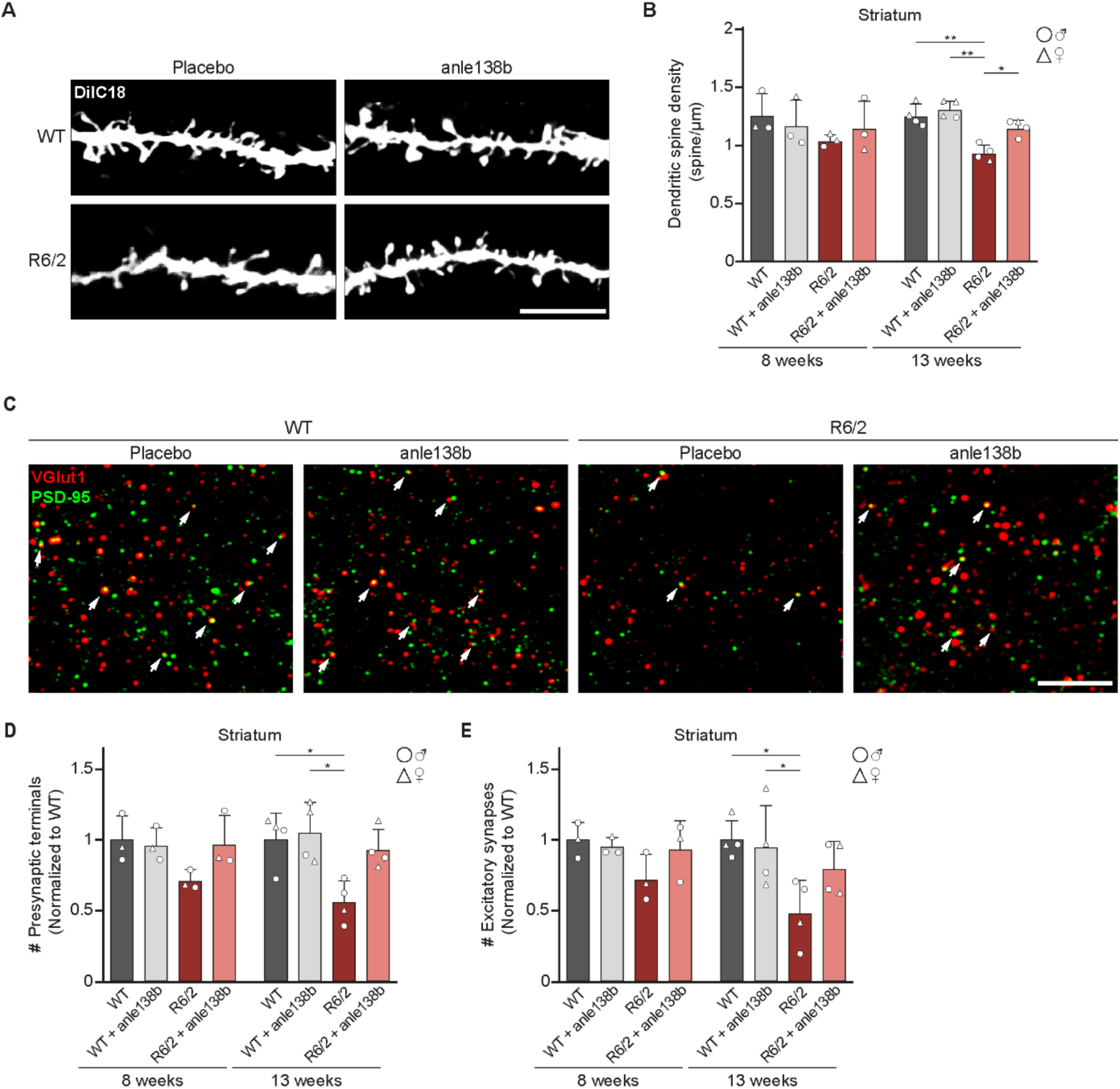
Anle138b mitigates synapse loss in the striatum of R6/2 mice. **(A)** Representative images of DiIC18 labeling showing dendrites with spines in the striatum of 13-week-old WT and R6/2 mice treated with placebo or anle138b. **(B)** Quantification of dendritic spine density in the striatum of 8- and 13-week-old WT and R6/2 mice. Two-way ANOVA with Bonferroni’s multiple comparison test, per age group. 8 weeks: not significant; 13 weeks: Treatment, **p < 0.0053; Genotype, ****p < 0.0001; Treatment x Genotype, p = 0.0517. n = 3 mice (8 weeks) or 4 mice (13 weeks) per genotype and treatment group. **(C)** Representative images of immunostained VGlut1 and PSD-95 puncta in the striatum of 13-week-old WT and R6/2 mice treated with placebo or anle138b. Excitatory synapses were identified by the overlap between the puncta in the two channels (examples indicated by white arrows). **(D)** Quantification of the number of VGlut1 puncta in the striatum of 8 and 13-week-old WT and R6/2 mice. Two-way ANOVA with Bonferroni’s multiple comparison test, per age group. 8 weeks: not significant; 13 weeks: Treatment, p = 0.07; Genotype, **p < 0.0059; Treatment x Genotype, p = 0.0688. n = 3 mice (8 weeks) or 4 mice (13 weeks) per genotype and treatment group. **(E)** Quantification of the number of overlapping VGlut1 / PSD-95 puncta in the striatum of 8 and 13-week-old WT and R6/2 mice. Two-way ANOVA with Bonferroni’s multiple comparison test, per age group. 8 weeks: not significant; 13 weeks: Treatment, p = 0.2670; Genotype, *p < 0.0122; Treatment x Genotype, p = 0.1217. n = 3 mice (8 weeks) or 4 mice (13 weeks) per genotype and treatment group. Data presented as mean ± SD. Significant pairwise comparisons are indicated on the graphs. *p < 0.05, **p < 0.01. Scale bars in A and C, 5 µm.

Dendritic spines are a major site of excitatory synaptic inputs. We therefore assessed the density of excitatory synapses by co-immunostaining for the excitatory presynaptic marker VGlut1 and postsynaptic marker PSD-95. We observed lower density of VGlut1-positive presynaptic terminals in placebo-treated 13-week-old R6/2 mice (Fig. 6C-D). Accordingly, the density of synapses assessed by the number of overlapping VGlut1 and PSD-95 puncta was also significantly reduced (Fig. 6E). In contrast, the density of presynaptic terminals and overlapping VGlut1 and PSD-95 puncta in anle138b-treated R6/2 mice was not significantly different from wildtype littermates (Fig. 6C-E). These findings suggest that anle138b partially prevents synapse loss in striatal medium spiny neurons of R6/2 mice.

### Anle138b ameliorates disease phenotypes in zQ175DN knock-in HD mice

Having observed a clear disease-modifying effect of anle138b in R6/2 mice, we sought to validate these findings in another HD model. To this end, we used heterozygous zQ175DN knock-in mice expressing full-length *Htt* with ∼190 CAG repeats from the endogenous murine *Htt* locus (Menalled et al., 2012; Southwell et al., 2016). As treatments during the entire pre- and postnatal development are difficult in human patients, we furthermore tested a different anle138b administration regime. The compound was delivered at the same concentration of 2 g/kg as in the R6/2 model, but the treatment was started in adult mice at 4 months of age, and the phenotypic analyses were conducted in 9-month-old animals (Fig. 7A). We did not observe clear behavioral defects in heterozygous zQ175DN mice in a previous study (Voelkl et al., 2023), therefore here we focused on the assessment of brain morphology, mHTT aggregation and expression of striatal markers.

**Figure 7.**
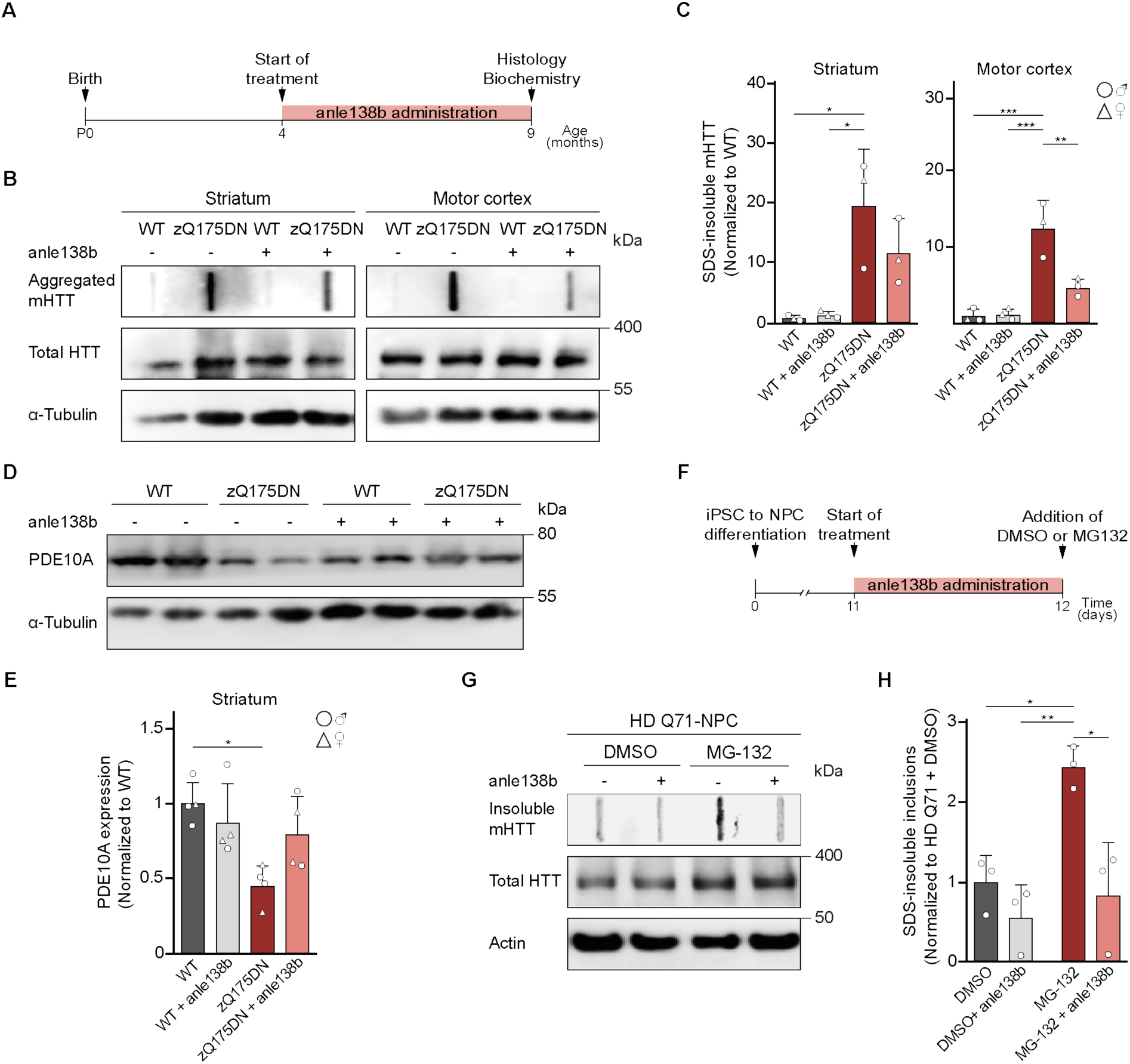
Anle138b mitigates HD phenotype in zQ175DN mice and rescues mHTT aggregation in human HD-iPSC derived NPCs. **(A)** Experimental timeline for the zQ175DN mice. **(B)** Filter trap of lysates from the striatum and motor cortex of 9-month-old WT and zQ175DN mice treated with placebo or anle138b. Total levels of HTT were determined by immunoblotting, α-Tubulin was used as loading control. **(C)** Quantification of aggregated mHTT in the striatum and motor cortex of 9-month-old WT and zQ175DN mice. Values were normalized to WT/placebo. Two-way ANOVA with Bonferroni’s multiple comparison test. Striatum: Treatment, p = 0.2715; Genotype, **p = 0.0017; Treatment x Genotype, p = 0.2208. Motor cortex: Treatment, **p = 0.01; Genotype, ***p = 0.0001; Treatment x Genotype, **p = 0.0074. n = 3 - 4 mice per group. **(D)** Representative immunoblot for PDE10A expression in striatal lysates of 9-month-old WT and zQ175DN mice treated with placebo or anle138b. α-Tubulin was used as loading control. **(E)** Quantification of PDE10A expression levels. Values were normalized to WT/placebo. Two-way ANOVA with Bonferroni’s multiple comparison test. ANOVA: Treatment, p = 0.3005; Genotype, **p = 0.01; Treatment x Genotype, *p = 0.0390. n = 3 - 4 mice per group. **(F)** Experimental timeline for HD-iPSC derived NPC cultures. **(G)** Filter trap of NPCs treated with vehicle (DMSO) or 7 µM anle138b. Total levels of HTT were determined by immunoblotting, Actin served as loading control. **(H)** Quantification of aggregated mHTT. Values were normalized to vehicle-treated NPCs. Two-way ANOVA with Bonferroni’s multiple comparison test. Anle138b, **p = 0.0041; MG-132, **p = 0.0094; anle138b x MG-132, *p = 0.0485. n = 3 independent experiments. Data presented as mean ± SD. Significant pairwise comparisons are indicated on the graphs. *p < 0.05, **p < 0.01, ***p < 0.001.

We did not detect significant differences in the area of the whole forebrain of zQ175DN mice compared to wildtype littermate controls, nor in the volume of the whole brain, the lateral ventricles, striatum or motor cortex (Appendix Fig. S3A-D). Immunostaining for GFAP and Iba1 also did not reveal signs of astro- or microgliosis at 9 months of age (Appendix Fig. S4A-D). We therefore could not assess potential beneficial effects of anle138b on these parameters. The fraction of neurons with mHTT inclusion bodies in the striatum and motor cortex was not different between placebo- and anle138b-treated zQ175DN animals (Appendix Fig. S5A-B). However, filter trap assay showed significant accumulation of aggregated mHTT species in the striatum and motor cortex of placebo-treated zQ175DN mice, and this accumulation was reduced by anle138b treatment in the motor cortex (Fig. 7B-C). We next quantified the protein levels of striatal markers by western blot. While the levels of DARPP-32 were not significantly changed in zQ175DN mice at 9 months of age (Appendix Fig. S5C-D), PDE10A levels were lower in placebo-treated, but not in anle138b-treated mice (Fig. 7D-E). Taken together, these findings demonstrate that anle138b reduces mHTT aggregation and prevents loss of PDE10A in the zQ175DN full-length knock-in model of HD.

### Anle138b reduces mHTT aggregates in human HD-iPSC-derived neural precursor cells

Having observed positive effects of anle138b in murine neurons and HD mouse models, we sought to validate these findings in a human model system. To this end, we differentiated iPSCs from a human HD patient into neural progenitor cells (NPCs) and treated the cultures for 24 hrs with anle138b or vehicle control. As NPCs do not show spontaneous mHTT inclusion formation, the proteasome inhibitor MG-132 was then applied to induce aggregation of mHTT inclusions (Fig. 7F). Filter trap analysis revealed a marked increase in aggregated mHTT in vehicle-treated cultures upon addition of MG-132, which was greatly reduced by anle138b (Fig. 7G-H). These results demonstrate the efficacy of anle138b in reducing mHTT aggregation in a human model system with endogenous expression of mHTT.

## Discussion

Here, we show that the small molecule anle138b significantly reduces mHTT toxicity and ameliorates disease phenotypes in a primary neuron model and two different mouse models of HD. In mice, consistent improvements were observed in all behavioral tests and multiple neuropathological and neurochemical disease signatures. The treatment had beneficial effects not only when administered throughout the embryonic development and postnatal life, but also when started at an adult age. Importantly, anle138b inhibited mHTT aggregation in human NPCs differentiated from HD-iPSCs, confirming its efficiency in human neuronal cells with endogenous full-length mHTT. Of note, anle138b did not alter the expression of wildtype HTT and did not cause adverse effects in wildtype mice in any of the described assays. These findings indicate that anle138b may be a possible therapeutic option for HD.

A number of other small molecules have shown disease-modifying potential in HD mice (Di Pardo et al., 2014; Ferrante et al., 2004; Hsiao et al., 2014; Masuda et al., 2008; Peng et al., 2008; Reick et al., 2016; Reiner et al., 2012; Squitieri et al., 2015). One of these compounds, the sigma-1 receptor agonist Pridopidine (Squitieri et al., 2015), is currently in clinical testing. Another compound, the vesicle monoamine transporter inhibitor Deutetrabenazine, is used for treating HD-associated chorea (Anderson et al., 2017). The splicing modulator Branaplam showed promising results in mouse and iPSC models of HD (Keller et al., 2022; Krach et al., 2022), but turned out to have neurotoxic side effects in a phase 2 clinical trial (NCT05111249), illustrating the challenges of developing small molecule therapeutics for HD. Of note, the mechanisms of action of these drugs are distinct from anle138, which reduces mHTT aggregate load without detectable impact on other cellular pathways. While the effects of anle138b in HD mice are similar in magnitude to other small molecules, many other treatments had to be administered through subcutaneous, intraperitoneal or intraventricular injections. In contrast, anle138b has excellent oral bioavailability. Moreover, the available results from clinical phase 1 trials indicate a favorable safety and tolerability profile in humans (Levin et al., 2022).

In both HD models used in the study, we administered anle138b to presymptomatic animals before the onset of prominent degenerative changes in the brain and manifestation of neurological symptoms. As HD is a hereditary disorder and mutation carriers can be identified in the presymptomatic phase, preventive approaches for this disease are feasible. In future studies it will be interesting to determine whether anle138b treatment is also beneficial when started after symptom onset.

It should be noted that while behavioral and neuropathological phenotypes were clearly improved by anle138b treatment in animals of either sex, a positive effect on life span was detected only in female mice. Sex-specific differences in clinical manifestations and treatment efficacy are documented in human patients and animal models of several neurodegenerative diseases including HD. In particular, females HD patients show on average more severe symptomology with regard to both motor and cognitive deficits (Hentosh et al., 2021). Furthermore, several experimental treatments for HD produced effects of different magnitude depending on the sex of the animals (Ma et al., 2007; Reiner et al., 2012), in line with our observations for anle138b. The causes of these differences remain poorly understood and represent an important subject for future research.

What could be the potential cellular substrate for anle138b-mediated improvements in neurological symptoms? We show that the treatment partially prevented loss of spines and excitatory synapses on striatal medium spiny neurons. A major source of excitatory inputs to the striatum is the neocortex, and dysfunction and degeneration of the corticostriatal afferents is believed to be a likely trigger of medium spiny neuron demise (Deng et al., 2013; Estrada-Sanchez and Rebec, 2013). Improved synaptic integrity and corticostriatal communication is therefore a likely contributing factor in the beneficial effects of anle138b, although other brain regions and cellular substrates might also play a role.

Further experiments are needed to clarify the mechanistic details of anle138 action at the molecular level. The rescue of HD-related phenotypes in all utilized models correlated with the ability of anle138b to reduce mHTT aggregate load, suggesting that the functional improvements likely result from a reduction in aggregated species of mHTT. Experiments with other purified recombinant aggregating proteins such as tau and α-synuclein showed that anle138b can directly modulate aggregation by binding to oligomers (Albariqi et al., 2024; Heras-Garvin et al., 2019; Martinez Hernandez et al., 2018; Wagner et al., 2015; Wagner et al., 2013; Wegrzynowicz et al., 2019). It should be kept in mind that in living cells, disease-related protein aggregates including mHTT aggregates show a great heterogeneity of conformations and biological effects (Gracia et al., 2020; Isas et al., 2021; Riguet et al., 2021;

Swanson et al., 2025). Actions of anle138b may therefore be distinct depending on the mHTT inclusion type. In the human brain, two major subcellular locations of mHTT inclusion bodies are the nucleus and the cytoplasm of neurites (DiFiglia et al., 1997). Anle138b reduced the frequency and toxicity of both nuclear and cytoplasmic inclusions in primary neurons. Interestingly, nuclear and cytoplasmic mHTT inclusions appear to have different protein composition, biochemical and ultrastructural properties, and distinct mechanisms of toxicity (Landles et al., 2020; Riguet et al., 2021). In particular, our previous findings indicate that cytoplasmic mHTT inclusions markedly impair cellular proteostasis (Blumenstock et al., 2021). Using the Fluc-EGFP proteostasis reporter, we show that anle138b ameliorates this impairment. Nuclear inclusions might be less burdensome for the proteostasis system, and the details of anle138b effects in this compartment remain to be explored. In summary, our results provide evidence of the promise of anle138b as a disease-modifying therapy for HD. As CAG trinucleotide repeat disorders share several common mechanisms (Everett and Wood, 2004), anle138b might also be beneficial in CAG expansion diseases beyond HD.

## Materials and Methods

### Plasmids

For transfection, the following plasmids were used: pcDNA3.1 mCherry, pcDNA3.1 HTT-Exon1-Q25-mCherry-myc-His and pcDNA3.1 HTT-Exon1-Q97-mCherry-myc-His (Hipp et al., 2012); pcDNA3.1 HTT-Exon1-Q25-His and pcDNA3.1 HTT-Exon1-Q72-His (Jeong et al., 2011; Voelkl et al., 2023); and NES-Fluc-EGFP (Park et al., 2013). The sequence of HTT-Exon1-Q25-mCherry-myc-His and HTT-Exon1-Q97-mCherry-myc-His and the sequence of HTT-Exon1-Q25-His and HTT-Exon1-Q72-His, are identical apart from the polyQ length.

### Mouse lines

All animal experiments were approved by the Government of Upper Bavaria (permit numbers ROB-55.2-2532.Vet_02-20-05 and ROB-55.2-2532.Vet_02-19-83) and conducted in accordance with the relevant guidelines and regulations. Mice were housed at the animal facility of the Max Planck Institute of Biochemistry under specific-pathogen-free conditions and *ad libitum* access to food and water. Transgenic R6/2 mice (Mangiarini et al., 1996) (JAX stock #002810) were maintained by breeding hemizygous R6/2 males with the F1 female progeny of the cross between CBA (Janvier Labs) and C57BL/6 (Janvier Labs) mice. Knock-in zQ175DN mice (Menalled et al., 2012; Southwell et al., 2016) (JAX stock #029928) were maintained on a C57BL/6 background. Anle138b treatment was administered orally, as food pellets (2 g/kg; Ssniff Spezialdiäten). Control groups were administered food pellets of the same composition, but without anle138b (placebo). For all experiments, mice of either sex were used, and experimenters were blind to the treatment groups. R6/2 CAG repeat tract length was determined from ear biopsies by Transnetyx® and amounted to 206 ± 10 repeats.

### Anle138b and BSA complexation

For application in an aqueous medium, a 10 mM stock solution of anle138b in dimethyl sulfoxide (DMSO) was prepared. Next, anle138b in DMSO was dissolved in an equimolar amount of bovine serum albumin (BSA; Roth) in phosphate buffer saline (PBS) to a final theoretical concentration of 200 µM. The final concentration of the anle138b/BSA complex was determined by high-performance liquid chromatography to be 165 µM.

### Primary neuronal cultures

Primary cortical neurons were prepared from E15.5 CD-1 wild type mouse embryos of either sex. Pregnant female mice were sacrificed by cervical dislocation, the uterus removed from the abdominal cavity and the embryos harvested and decapitated in a sterile Petri dish with ice-cold dissection medium, consisting of Hanks’ balanced salt solution (HBSS; Invitrogen) supplemented with 0.01 M HEPES, pH 7.4, 0.01 M MgSO_4_ and 1 % penicillin/streptomycin (Invitrogen). The brain was removed from the skull, the hemispheres separated, the meninges removed and the cortices isolated. The individual cortices were digested with 0.25 % trypsin containing 1 mM ethylenediaminetetraacetic acid (EDTA) and 15 μl 0.1 % DNAse I for 20 min at 37 °C. The digestion was stopped by washing with Neurobasal medium (Invitrogen) containing 5 % fetal bovine serum, and cells were dissociated by triturating the tissue in pre-warmed Neurobasal medium. Cells were centrifuged at 130 x g for 5 min, the supernatant was removed, and the pellet resuspended in Neurobasal medium supplemented with 2 % B27 (Invitrogen), 1 % L-Glutamine (Invitrogen) and 1 % penicillin/streptomycin. The neurons were then plated in 24-well plates on 13 mm sterile coverslips coated with 50 µg/ml poly-D-Lysine (Sigma) and 1 µg/ml laminin (ThermoFisher Scientific) at a density of 120,000 neurons per coverslip.

### Transfection of primary neurons

Neurons were transfected at day *in vitro* (DIV) 7 using the NeuroMag Transfection Reagent (OZ Biosciences), following the manufacturer’s instructions. Briefly, the transfection solution was prepared by adding 1 µg of NeuroMag reagent per µg of DNA, to 50 µL of pre-warmed Neurobasal medium containing 1 µg of DNA per construct. The mixture was incubated at room temperature (RT) for 20 min and added to the neurons in a 24-well plate. The cultures were placed on a magnetic plate at 37 _°_C, 5% CO_2_ for 15 min. Finally, the magnetic plate was removed and the neurons kept at 37 _°_ C, 5 % CO_2_ for protein expression.

### Anle138b treatment and proteasome inhibition in primary neurons

Neurons were treated at DIV 7 + 1 with 7 µM of anle138b/BSA in PBS:DMSO (3:1) or DMSO as control, for 24 hr at 37 _°_C, 5 % CO_2_. For proteasome inhibition, neurons were treated at DIV 7 + 2 with 5 µM of MG-132 or DMSO as control, for 4 hr at 37_°_ C, 5 % CO_2_.

### Immunocytochemistry on primary neurons

Transfected neurons were fixed at DIV 7+2 with 4 % paraformaldehyde (PFA; Electron Microscopy Sciences) in PBS for 15 min at RT. Remaining free PFA groups were quenched with 50 mM ammonium chloride in PBS for 10 min at RT. Neurons were rinsed once with PBS, permeabilized with 0.25 % Triton X-100 (Merck) in PBS for 5 min and washed 3 x 5 min with PBS. Following the washes with PBS, neurons were incubated in blocking solution consisting of 2 % BSA, 4 % donkey serum (Jackson Immunoresearch) and 0.01 % NaN_3_ in PBS for 30 min at RT. After blocking, coverslips were transferred to a light protected humid chamber and incubated with primary antibodies diluted in blocking solution for 1 hr. The following primary antibodies were used: goat anti-mCherry (1:300; AB0040-200, Origene), rabbit anti-cleaved caspase-3 (1:400; 9662, Cell Signaling Technologies), mouse anti-EM48 (1:300; MAB5374, Merck), chicken anti-MAP2 (1:500; NB300-213, Novus Biological) and mouse anti-His (1:300; ab18184, Abcam). Neurons were washed 3 × 10 min with PBS at RT and incubated for 30 min with Alexa Fluor 488, Alexa Fluor 647 and Cyanine Cy3 conjugated secondary antibodies, derived from donkey, (1:300; Jackson Immunoresearch) and 0.5 µg/ml DAPI in blocking solution. Neurons were washed 2 × 10 min with PBS, rinsed once with Mili-Q water and mounted on Menzer glass slides (VWR) using Prolong Glass Antifade mounting medium (Invitrogen).

Images were acquired using a Thunder Imager 3D Tissue fluorescent microscope (Leica) and analyzed using the ImageJ/Fiji software (Schindelin et al., 2012). For neuronal viability analysis, neurons with a negative cleaved caspase-3 signal were considered to be viable. For the analysis of Fluc-EGFP foci containing cells, a value of 2 standard deviations above the foci/cytoplasm intensity ratio was considered as the threshold for the presence of foci. Whenever possible, the experimenter was blind to the treatment groups.

### Behavioral tests and life span analysis

All behavioral analyses were performed during the mice’s dark phase and habituation to the testing room was done for a period of 30 – 60 min prior to any testing. For the grip strength test, a total of three trials were performed per mouse. For each trial, forelimb strength was measured with the bar version of the BIO-GS3 Grip Test (Bioseb). The final score was the average of the three trials. Rotarod performance was analyzed using the Rota-Rod NG system (Ugo Basile) and it consisted of two training days followed by a test day. During training the mice were placed on the rotating rod at 5 rotations per minute (rpm) for 5 min. For the test, a speed gradient of 5 – 40 rpm was implemented and the latency to fall was measured up to a limit of 5 min. Three trials were performed, and the results averaged to obtain the final score. The open field test consisted of a single trial in which mice were recorded in an open arena (40 x 40 x 40 cm) with black walls, white floor and illuminated homogeneously from above. Mice were recorded for 15 min and distance travelled was calculated automatically using the EthoVision XT 16 software (Noldus Information Technology). For limb clasping, mice were recorded for 20 seconds while held by their tail, approximately 30 cm above a bench top. Limb clasping was assessed on three consecutive days and the time spent clasping with either fore-, hind- or fore- and hindlimbs was manually recorded. The final value was the average of the three trials. Life span was determined by monitoring mouse burden according to score sheet based on appearance, body weight and rearing reflex on a daily basis. Mice with severe burden were considered to have reached the experimental endpoint.

### Immunohistochemistry

Mice were deeply anesthetized with 1.6 % ketamine/0.08 % xylazine and transcardially perfused with PBS followed by 4 % PFA in PBS. Brains were extracted from the skull and post-fixed in 4 % PFA in PBS overnight at 4 _°_C. Fixed brains were sectioned into 30 µm thick coronal sections using a VT1000S vibratome (Leica). Floating sections were permeabilized with 0.5 % Triton X-100 in PBS for 30 min at RT. Following permeabilization, staining against the VGlut1 and PSD95 markers required antigen retrieval with 20 µg/ml Proteinase K (Sigma-Aldrich) in 50 mM Tris Base, 1 mM EDTA and 0.5 % Triton X-100 in H_2_O for 10 min at 37 _°_ C. Sections were washed once with PBS for 10 min at RT and incubated in blocking solution consisting of 0.2 % BSA, 5 % donkey serum, 0.2 % L-Lysine (Sigma-Aldrich), 0.2 % Glycine (Sigma-Aldrich) and 0.01 % NaN_3_ in PBS for 1.5 hr at RT. After blocking, sections were incubated with primary antibodies diluted in 2 % BSA, 0.3 % Triton X-100 and 0.01 % NaN_3_ in PBS at 4 _°_C overnight. The following primary antibodies were used: mouse EM48 (1:300; MAB5374, Merck), chicken anti-MAP2 (1:500; NB300-213, Novus Biological), mouse anti-NeuN (1:400; MA5-33103, ThermoFisher Scientific), rabbit anti-DARPP-32 (1:400; ab40801, Abcam), chicken anti-GFAP (1:500; AP31806PU-N, Origene), goat anti-Iba1 (1:300, ab107159, Abcam), mouse anti-VGlut1 (1:300; ab134283, Abcam) and rabbit anti-PSD95 (1:300; GTX133091, GeneTex). Sections were washed 3 × 10 min with PBS at RT and incubated for 1 hr at RT with Alexa Fluor 488, Cyanine Cy3 and/or Alexa Fluor 647 conjugated secondary antibodies, derived from donkey (1:300; Jackson Immunoresearch), NeuroTrace 640/660 (1:300; ThermoFisher Scientific) and 0.5 µg/ml DAPI in 0.3 % Triton X-100, 3 % donkey serum and 0.01 % NaN_3_ in PBS. Sections were washed 3 × 10 min with PBS and mounted on Menzer glass slides using Prolong Glass Antifade mounting medium.

Images were acquired using a STELLARIS 5 confocal microscope (Leica) and analyzed using the ImageJ/Fiji software. For the quantification of forebrain area, a region of interest (ROI) was manually drawn around the forebrains. The volume of the striatum, motor cortex, ventricles and whole brain was analyzed according to the principle of Cavalieri, as previously described (Cyr et al., 2005) (volume = s_1_d_1_ + s_2_d_2_ + … s_n_d_n_, where s = surface area and d = distance between sections). For volume estimation, 9 sections spaced 200 µm apart were used. The boundaries of the individual brain regions were defined by anatomical landmarks identifiable by NeuN staining according to the Allen Brain Atlas (mouse.brain-map.org). Immunoreactivity was analyzed in the striatum and motor cortex by identifying these areas, manually drawing a ROI and measuring their size. Background correction was used when necessary and the area occupied by the appropriate immunofluorescent signal (GFAP or Iba1) was quantified by adequate thresholding and morphological filtering. The percentage of ROI area occupied by immunoreactive species was determined by calculating the ratio between the two areas. The percentage of inclusion bearing neurons was analyzed in the striatum and motor cortex by counting the mHTT inclusions overlapping with nuclear DAPI signal. For the quatification of excitatory synapses, images of pre- and postsynaptic immunostainings (VGlut1 and PSD95, respectively) were superimposed using the ImageJ “Colocalization Threshold” feature. Puncta smaller or bigger than 2 or 10 pixels, respectively, and puncta with a circularity smaller than 0.8 were excluded from the analysis. The resulting colocalization map was isolated and the and the number of puncta quantified using the ImageJ “Analyze Particles” feature. Whenever possible, the experimenter was blind to the treatment groups.

### Dendritic spine labeling

Fixed brain sections were labelled with the lipophilic dye 1,1’-Dioctadecyl-3,3,3’,3’-tetramethylindocarbocyanine perchlorate (DiIC18; ThermoFisher Scientific) as previously described (Speranza *et al*., 2022). Briefly, sections were placed on a glass slide, the PBS was removed and DiIC18 crystals were applied directly into the striatum. For this, the tip of a 27-gauge needle was coated with DiIC18 and the crystals slowly deposited in the regions of interest, taking care to avoid excessive amounts of crystals in the same area. The sections were covered with PBS to avoid dehydration and incubated in the dark for 15 min, at RT. Next, the sections were transferred to a 24-well plate and incubated in PBS at 4 _°_ C, protected from light, for 3 o/n. For combined immunofluorescence, the sections were permeabilized with 100 µg/mL Saponin (ThermoFisher Scientific), 3 % BSA in PBS at RT, for 30 min. After permeabilization, sections were incubated with primary antibodies diluted in 100 µg/mL Saponin, 3 % BSA and 0.01 % NaN_3_ in PBS at 4 _°_ C, o/n. The following primary antibodies were used: chicken anti-MAP2 (1:500; NB300-213, Novus Biological) and rabbit anti-DARPP-32 (1:400; ab40801, Abcam). Sections were washed 3 × 10 min with PBS at RT and incubated for 3 hr at RT with Alexa Fluor 488 and Alexa Fluor 647 conjugated secondary antibodies, derived from donkey, (1:300; Jackson Immunoresearch) and 0.5 µg/ml DAPI in 3 % BSA and 0.01 % NaN_3_ in PBS. Finally, sections were washed 3 × 10 min with PBS and mounted on Menzer glass slides using Prolong Glass Antifade mounting medium.

Images were acquired using a STELLARIS 5 confocal microscope and analyzed using the ImageJ/Fiji software. For dendritic spine density analysis, a ROI was manually drawn around DiIC18 labelled dendrites, and their length measured. All lateral protrusions in the ROI were semi-automatically detected using the ImageJ plugin “Dendritic Spine Counter” and spine density expressed as the number of spines per dendritic length (n/µm). Whenever possible, the experimenter was blind to the treatment groups.

### Western blot and filter trap assay of brain tissue lysates

Mice were sacrificed by cervical dislocation, brains extracted from the skull and the striatum and motor cortex dissected on ice. Brain regions were homogenized on ice in lysis buffer (50 mM Tris buffer pH 7.4, 150 mM NaCl, 2 mM EDTA, 1 % Triton X-100, phosphatase inhibitor (Roche) and protease inhibitor cocktail (Roche)). The resulting lysates were centrifuged at 15,000 x g for 15 min at 4 _°_C, the pellet discarded, and total protein concentration was determined using the Protein Dye Assay kit (Bio-Rad). For Western blots, samples containing 20 µg of total protein were denatured at 95 _°_C for 5 min and separated according to their size on 10 % SDS-PAGE gels. The proteins were transferred onto polyvinylidene fluoride (PVDF) membranes (Bio-Rad), using a Trans-Blot Turbo transfer system (Bio-Rad), and unspecific binding sites were blocked by incubating the membranes in 5 % dried milk (Milipore), 3 % BSA in 0.1 % Tris-buffered saline with 0.1 % Tween 20 (ITW Reagents) (TBS-T) for 2 hrs at RT. After blocking, the membranes were incubated with primary antibodies diluted in 3 % BSA and 0.01 % NaN_3_ in TBS-T, overnight at 4 _°_C. The following primary antibodies were used: rabbit anti-PDE10A (1:1,000; ab227829, Abcam), rabbit anti-DARPP-32 (1:800; ab40801, Abcam), rabbit anti-HTT (1:1,000; 5656S, Cell Signaling Technologies), and mouse anti-α-Tubulin (1:2,000; T9026, Sigma Aldrich). Membranes were washed 3 × 10 min in TBS-T at RT and incubated for 2 hr at RT with horse-radish peroxidase (HRP) conjugated secondary antibodies, derived from donkey (1:2,500; ThermoFisher Scientific), diluted in 3 % dried milk and 0.01 % NaN_3_ in TBS-T. Proteins were visualized by chemiluminescence by adding the HRP substrate Clarity Max^TM^ Western ECL Substrate (Bio-Rad) to the membranes. Images were acquired using the FUSION FX system (Vilber).

For filter trap assays, tissue samples were centrifuged at 2,000 x g for 10 min at 4 _°_C, the pellet discarded, and total protein concentration calculated. Filter trap assays were performed with 35 µg of total protein diluted in 100 µL of lysis buffer. Cellulose acetate membranes (0.2 µm pore size, Bio-Rad) and filter paper (Bio-Rad) were pre-equilibrated in 0.1 % SDS in H_2_O for 10 min at RT and assembled in the slot blot apparatus (Bio-Dot SF, Bio-Rad). Samples were loaded and allowed to completely pass through the membrane under vacuum. Wells were washed 3 x with 100 µL 0.1 % SDS in H_2_O followed by standard immunoblotting of the membranes. For the detection of aggregated mHTT, mouse EM48 (1:1,000; MAB5374, Merck) primary antibody and HRP conjugated secondary antibody (1:2,500; ThermoFisher Scientific) were used.

### iPSC culture and differentiation into NPCs

The HD Q71-iPSC line was kindly provided by George Q. Daley. The HD Q71-iPSC line was established and characterized for pluripotency (Park et al., 2008). iPSCs were maintained on Geltrex (ThermoFisher Scientific) using mTeSR1 media (Stem Cell Technologies) at 37 °C, 5 % CO_2_. The absence of mycoplasma was confirmed by testing the iPSC lines for mycoplasma contamination at least once every 2 weeks. Neural differentiation was induced with STEMdiff Neural Induction Medium (Stem Cell Technologies) after the monolayer culture method (Chambers *et al*., 2009). In brief, iPSCs were washed once with PBS followed by the addition of 1 ml of Gentle Dissociation Reagent (Stem Cell Technologies). After 10 min incubation, the cells were gently collected and 2 ml of DMEM/F12 (ThermoFisher Scientific) containing 10 μM ROCK inhibitor (Abcam) was added. The cells were then centrifuged at 300 x g for 10 min and resuspended in STEMdiff Neural Induction Medium supplemented with 10 μM ROCK inhibitor. Cells were seeded on plates coated with 15 μg/ml poly-ornithine and 10 μg/ml laminin at a density of 200,000 cells/cm^2^ for neural differentiation.

### Anle138b treatment and proteasome inhibition in NPCs

NPCs were treated with 7 μM anle138b/BSA for 16 h. The following day, cells were treated with fresh 7 μM anle138b/BSA for 8 h in the presence of 5 μM MG-132 for proteasome inhibition or DMSO as control treatment.

### Western blot and filter trap assay in NPCs

Cells were collected and lysed in a non-denaturing lysis buffer (50 mM Hepes pH 7.4, 150 mM NaCl, 1 mM EDTA, 1% Triton X-100) supplemented with 2 mM sodium orthovanadate, 1 mM phenylmethylsulfonyl fluoride, and protease inhibitor mix on ice. Lysates were homogenized through syringe needle (27-gauge). Standard BCA protein assay was used to determine protein concentration. After equilibration of protein concentration, the equilibrated whole lysates were centrifuged at 8,000 x g for 5 min at 4 °C. Supernatant and pellet were separated for SDS-PAGE and filter trap analysis. The supernatant was used for SDS-PAGE analysis, transferred to PVDF membranes (Millipore), and subjected to immunoblotting. The following antibodies were used: anti-β-Actin (1:5,000; 8226, Abcam) and anti-HTT (1:1,000; ab5656, Cell Signaling Technologies). For the filter trap, the pellets were resuspended in 2 % SDS and loaded onto a cellulose acetate membrane assembled in a slot blot apparatus. The cellulose acetate membrane was rinsed with 0.2 % SDS and retained SDS-insoluble mHTT aggregates were detected with mouse anti-polyQ antibody (1:5,000; MAB1574, Millipore).

### Statistical analysis

Statistical analysis and graphical representations were performed in GraphPad Prism^TM^ v 10.2.3 (GraphPad Software Inc.). Bar graphs show the mean and standard deviation. Violin plots show the median, density curves and interquartile range. The significance level was set to p < 0.05, the statistical tests and group numbers are detailed in the figure legends.

## Acknowledgements

We thank Kerstin Voelkl for initial experiments at early stages of the project; Dennis Feigenbutz and Laura Dias for help with primary neuronal cultures; Laura Dias for help with brain volume quantification; Andreu Boix Pagès for assistance with striatal marker quantification; Yasmin Richter and Janine Kirstein for helpful input; Henry Klein and Elke Franke for excellent technical assistance; Magdalena Boehm for mouse genotyping and help with colony management. This work was funded by the Max Planck Society for the Advancement of Science and the Deutsche Forschungsgemeinschaft (DFG, German Research Foundation) - SPP2453 project number 541742535 (to I.D.), SFB1451 project number 431549029-A09 (to I.D.), CECAD, EXC 2030-390661388 (to D.V. and I.D) and EXC 2067/1-390729940 (to C.G.).

## Author contributions

M.S.P. performed experiments with primary neuronal cultures and mice, analyzed data and designed figures. S.K. conducted experiments with NPCs, which were supervised by D.V. E.C. performed dendritic spine labeling. S.R. provided reagents. A.G., C.G. and I.D. conceived the project. M.S.P., S.R., A.L., A.G., C.G. and I.D. interpreted the data. R.K. and I.D. supervised the project. M.S.P. and I.D. wrote the paper. M.S.P., D.V., A.G., C.G. and I.D. edited the manuscript with input from all the authors.

## Appendix figures

**Appendix figure S1.**
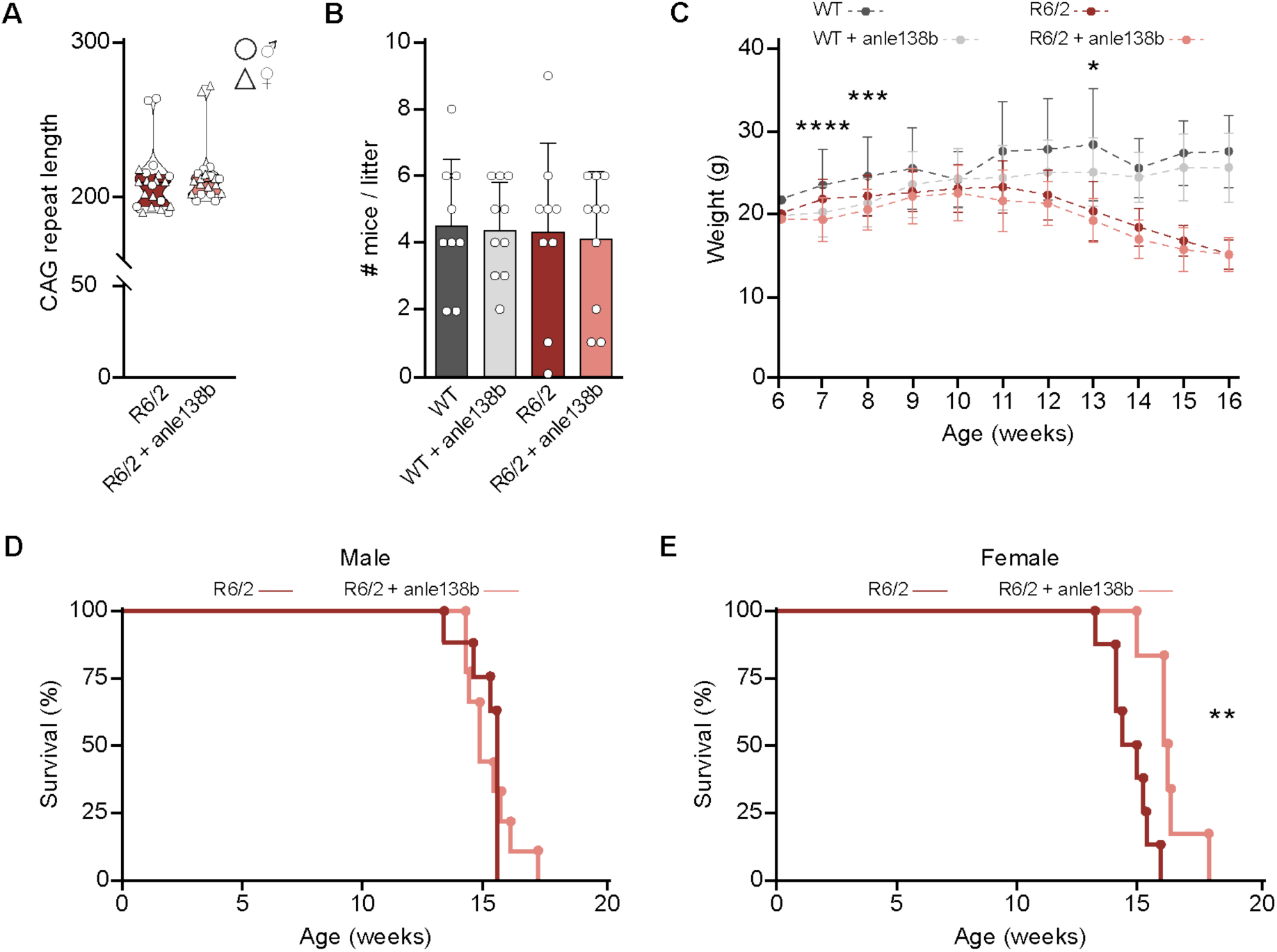
Additional characterization of anle138b-treated R6/2 mice. **(A)** CAG repeat length in 13-week-old R6/2 mice. Unpaired two-tailed *t*-test, not significant. n = 21 – 22 mice per treatment group. **(B)** Number of WT and R6/2 pups per litter for placebo or anle138b-treated breeding pairs. Two-way ANOVA, not significant. n = 9 – 10 litters per group. **(C)** Body weight of WT and R6/2 mice treated with placebo or anle138b. Repeated measures ANOVA. Significant differences between treatment groups are indicated on the graph for the respective time points: ***p<0.001; ****p<0.0001. n = 19 – 22 mice per group. **(D – E)** Kaplan-Meier survival curve for male D and female E R6/2 mice. Log-rank test. Significant pairwise comparisons are indicated on the graphs. **p < 0.01.

**Appendix figure S2.**
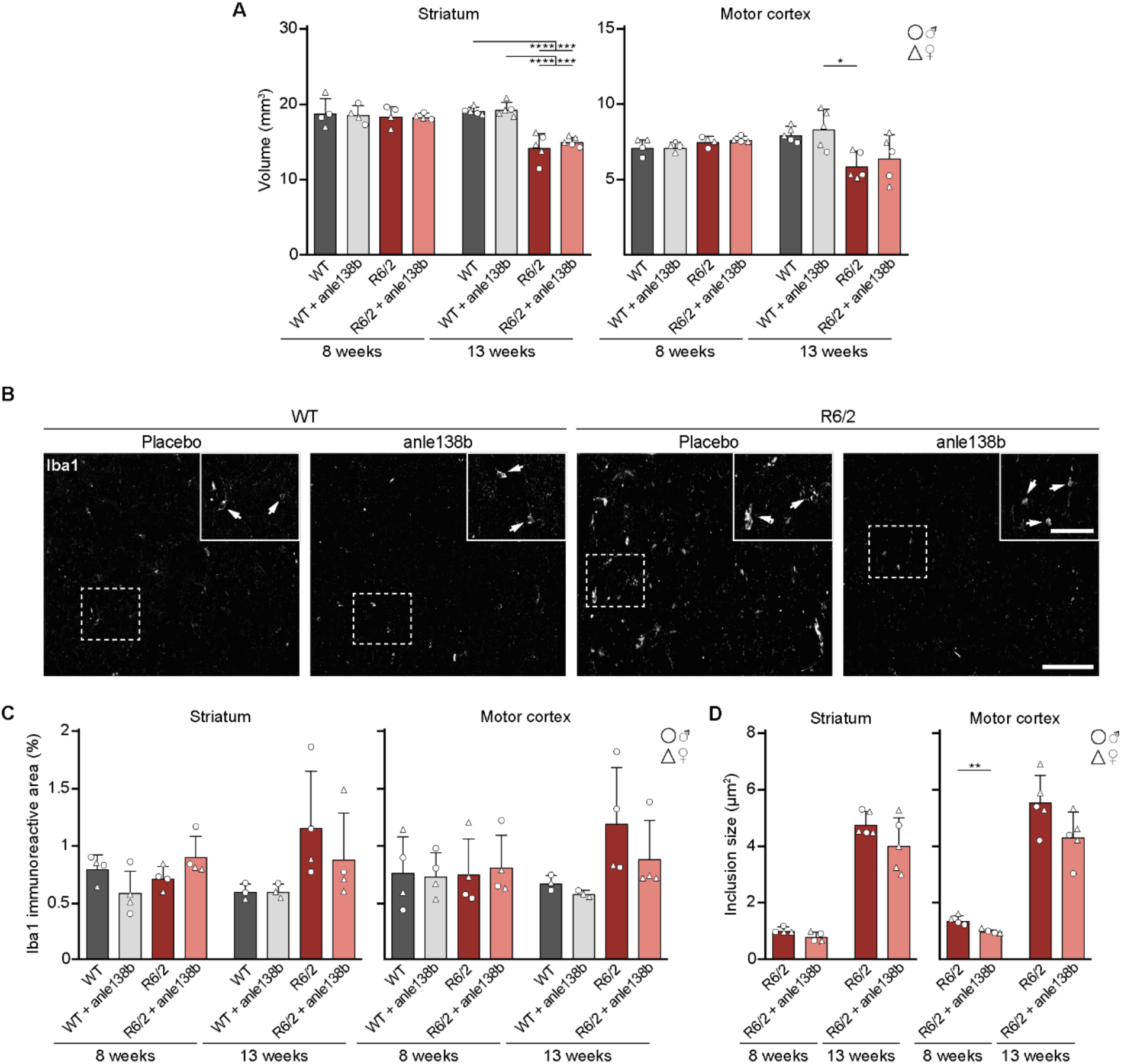
mHTT inclusion size and microgliosis in anle138b-treated R6/2 mice. **(A)** Striatum and motor cortex volume quantification in 8 and 13-week-old WT and R6/2 mice. Two-way ANOVA with Bonferroni’s multiple comparison test. Striatum, 8 weeks: not significant; 13 weeks: Treatment, p = 0.3682; Genotype, ****p < 0.0001; Treatment x Genotype, p = 0.5531. Motor cortex, 8 weeks: not significant; 13 weeks: Treatment, p = 0.4037; Genotype, **p < 0.0012; Treatment x Genotype, p = 0.9394. n = 4 (8 weeks) or 5 mice (13 weeks) per group. **(B)** Representative images of Iba1 immunostaining in the dorsal striatum of 13-week-old WT and R6/2 mice treated with placebo or anle138b. Insets show higher magnification of the areas delineated by the dashed boxes. White arrows point to examples of microglia. **(C)** Fraction of Iba1 immunopositive area in the striatum and motor cortex. Two-way ANOVA with Bonferroni’s multiple comparison test, not significant. n = 4 (8 weeks) or 3 – 4 mice (13 weeks) per group. **(D)** Quantification of mHTT inclusion size in the striatum and motor cortex of 8 and 13-week-old R6/2 mice. Unpaired two-tailed *t-*test, per age group and brain region. n = 4 (8 weeks) or 5 mice (13 weeks) per group. Data presented as mean ± SD. Significant pairwise comparisons are indicated on the graphs. *p < 0.05, **p < 0.01, ***p < 0.001, ****p < 0.0001. Scale bars in B, 100 µm; insets, 50 µm.

**Appendix figure S3.**
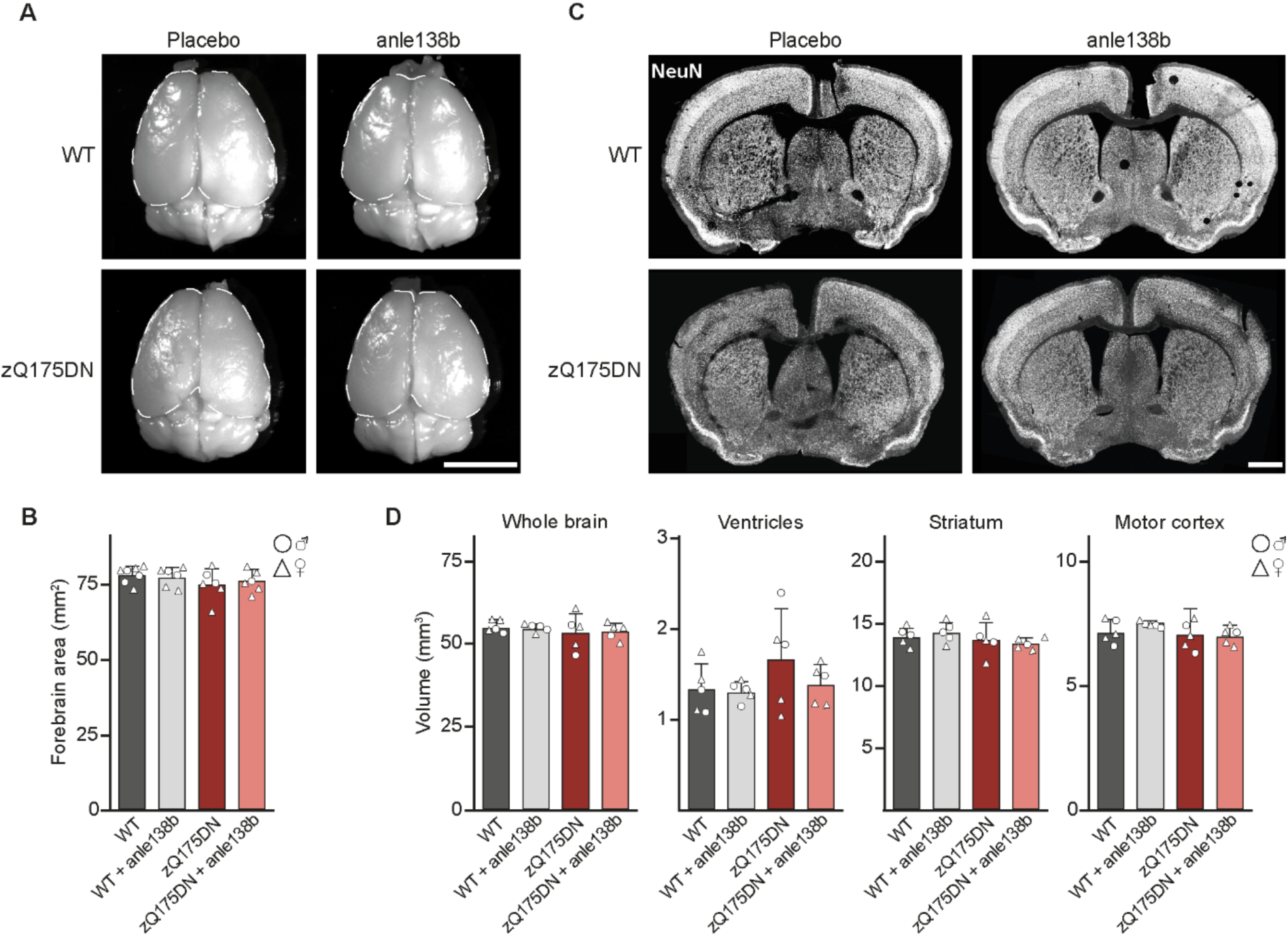
Effect of anle138b on brain morphology and mHTT inclusion load in zQ175DN mice. **(A)** Representative images of the brains of 9-month-old WT and zQ175DN mice treated with placebo or anle138b. Dashed white lines outline the forebrain. **(B)** Quantification of forebrain area in 9-month-old WT and zQ175DN mice. Two-way ANOVA with Bonferroni’s multiple comparison test, not significant. n = 6 – 7 mice per group. **(C)** Representative coronal brain sections of 9-month-old WT and zQ175DN mice treated with placebo or anle138b, immunostained for the neuronal marker NeuN. **(D)** Whole brain, ventricle, striatum and motor cortex volume quantification in 9-month-old WT and zQ175DN mice. Two-way ANOVA with Bonferroni’s multiple comparison test, not significant. n = 5 mice per group. Data presented as mean ± SD. Scale bars: A, 5 mm; C, 1 mm.

**Appendix figure S4.**
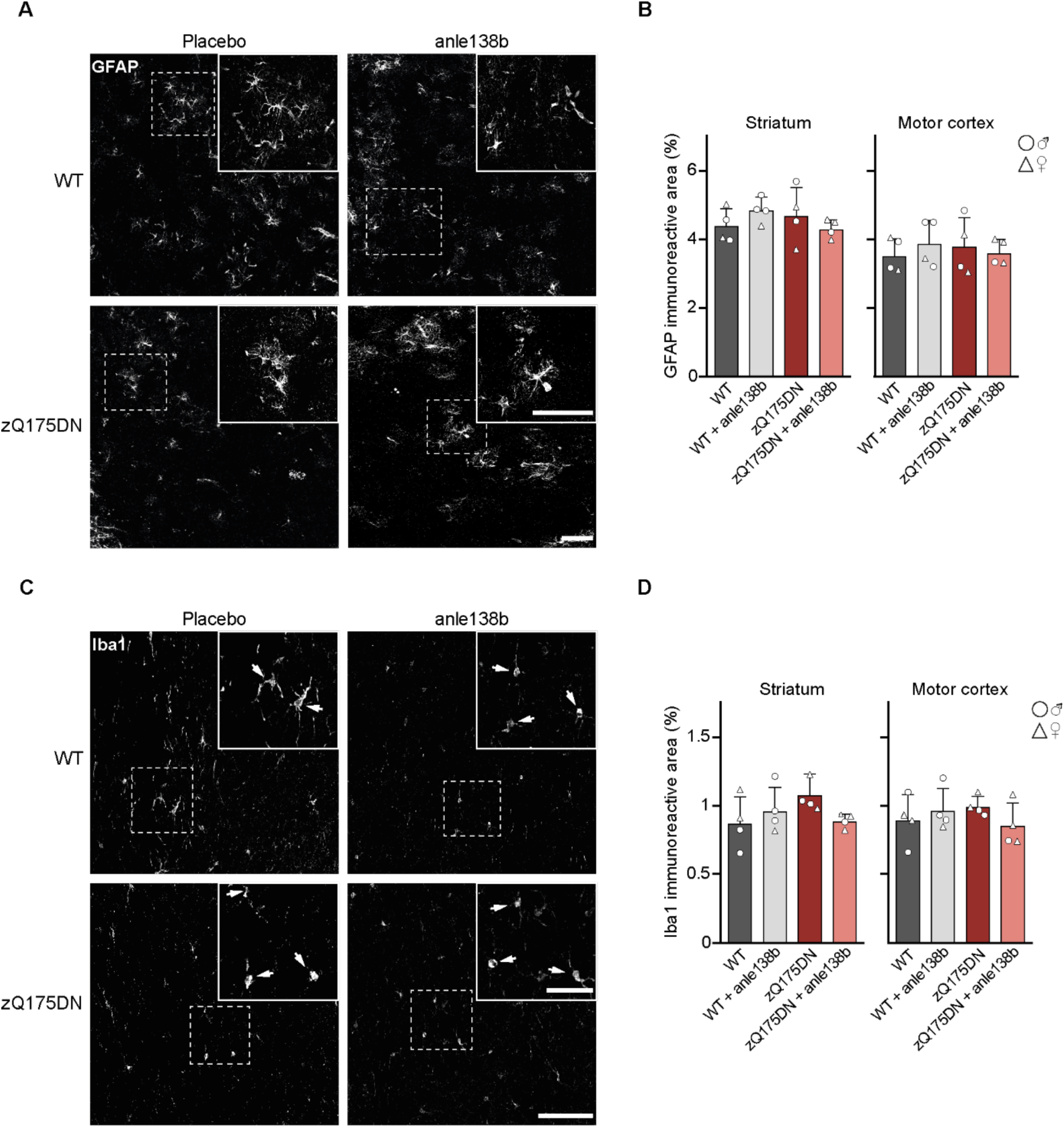
zQ175DN mice do not exhibit signs of neuroinflammation. **(A)** Representative images of GFAP immunostaining in the dorsal striatum of 9-month-old WT and zQ175DN mice treated with placebo or anle138b. Insets show higher magnification of the areas delineated by the dashed boxes. **(B)** Fraction of GFAP immunopositive area in the striatum and motor cortex of 9-month-old WT and zQ175DN mice. Two-way ANOVA with Bonferroni’s multiple comparison test, per brain region, not significant. **c** Representative images of Iba1 immunostaining in the dorsal striatum of 9-month-old WT and zQ175DN mice treated with placebo or anle138b. Insets show higher magnification of the areas delineated by the dashed boxes. Arrows point to examples of microglia. **(D)** Fraction of Iba1 immunopositive area in the striatum and motor cortex of 9-month-old WT and zQ175DN mice. Two-way ANOVA with Bonferroni’s multiple comparison test, per brain region, not significant. Data presented as mean ± SD. Scale bars: A and C, 100 µm; insets, 50 µm.

**Appendix figure S5.**
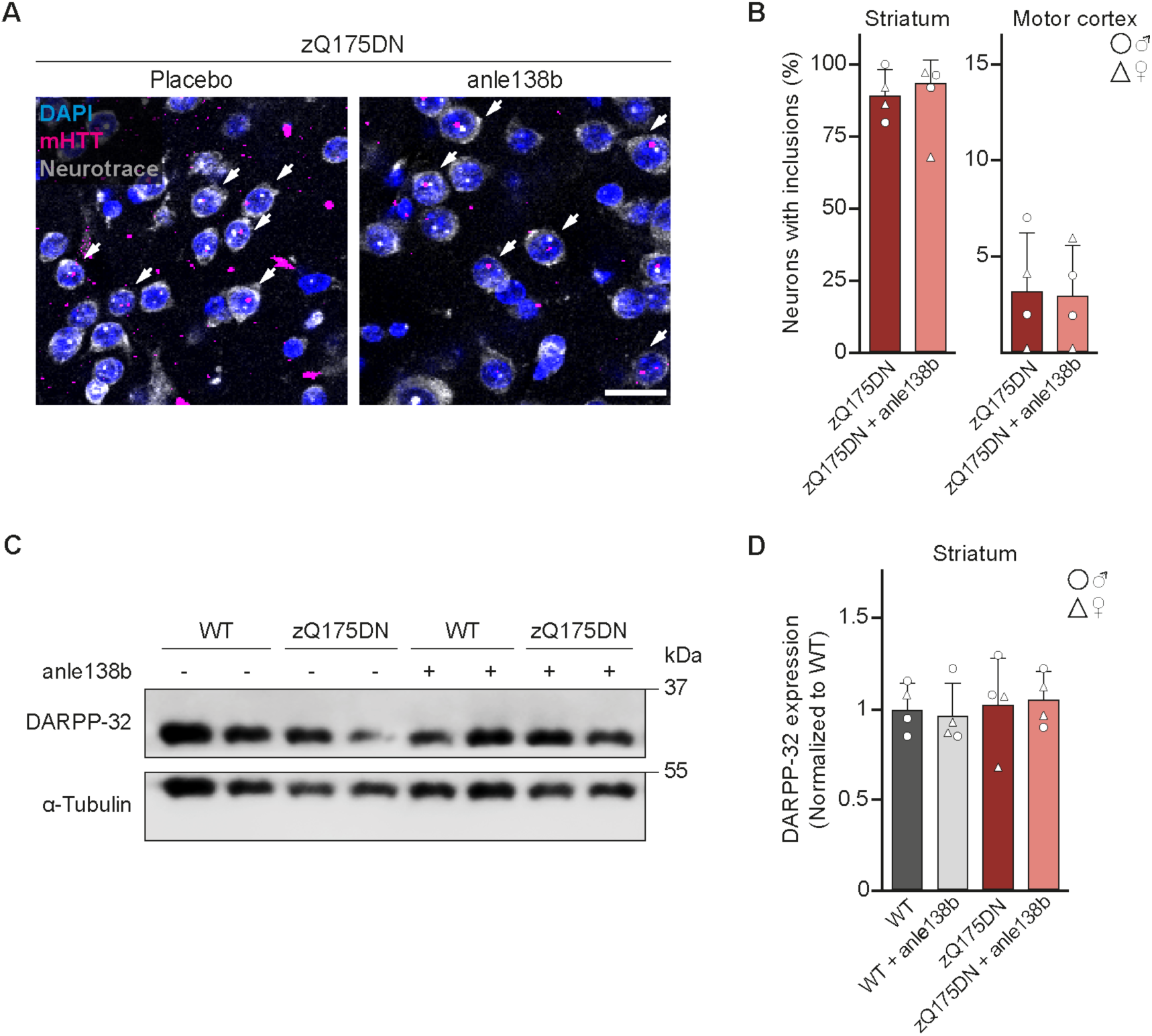
Anle138b does not alter inclusion load and DARPP32 expression in zQ175DN mice. **(A)** Representative neurons with nuclear mHTT inclusions (magenta) in the motor cortex of 9-month-old zQ175DN mice treated with placebo or anle138b. Neurons were identified by Neurotrace labeling and aggregated mHTT was detected by EM48 immunostaining. Nuclei were labelled with DAPI. White arrows point to neurons with mHTT inclusion bodies. **(B)** Quantification of the fraction of neurons with mHTT inclusion bodies in the striatum and motor cortex of 9-month-old zQ175DN mice. Unpaired two-tailed *t* test, not significant. n = 4 mice per treatment group. **(C)** Representative immunoblot for DARPP-32 in striatal lysates of 9-month-old WT and zQ175DN mice treated with placebo or anle138b. **(D)** Quantification of DARPP-32 expression levels. Values were normalized to WT/placebo. Two-way ANOVA with Bonferroni’s multiple comparison test, not significant. n = 4 mice per genotype and treatment group. Data presented as mean ± SD. Scale bar: A, 20 µm.

